# The tail domain of neurofilament light chain accumulates in neuronal nuclei during oxidative injury

**DOI:** 10.1101/2022.03.03.481279

**Authors:** Aleksandra Arsić, Ivana Nikić-Spiegel

## Abstract

Neurofilament light chain (NFL), one of the major subunits of neuron-specific intermediate filaments, is an important component of the neuronal cytoskeleton. Mutations in the *NEFL* gene and disturbances of NFL turnover or transport are associated with the pathophysiology of various diseases of the nervous system. Also, NFL is released from neurons as a result of neuroaxonal damage and NFL levels in cerebrospinal fluid and blood have emerged as a biomarker of the progression of numerous neurological diseases. One common underlying factor in these diseases is oxidative stress. Although both NFL and oxidative injury play a role in the development of neurological diseases, the effect of oxidative stress on NFL at the cellular level has not been extensively studied. Here, by investigating the response of neurofilaments to *in vitro* oxidative injury, we discovered that NFL localizes in the nuclei of primary neurons. We show that this depends on calpain activity and determine that a proteolytic fragment of the tail domain, not the full-length NFL, is that which accumulates in the nuclei. Furthermore, we demonstrate that the recombinant NFL tail domain accumulates in the nuclei of neurons and neuroblastoma cells, even in the absence of oxidative injury. Finally, we show that the tail domain of NFL interacts with the DNA in the nuclei of living cells, suggesting that NFL plays a role in the regulation of gene expression in response to oxidative injury.

## Introduction

Neurofilaments are neuron-specific cytoskeletal elements from the family of intermediate filaments^1,2^. They are heteropolymers composed of neurofilament light (NFL), medium (NFM), and heavy (NFH) chain subunits, and α-internexin in the central nervous system or peripherin in the peripheral nervous system^3-6^. These subunits interact to form long unbranched intermediate filaments with a diameter of 10 nm. Neurofilaments are sparse in neuronal cell bodies, dendrites, and synapses, but are highly abundant in axons where they form long, straight, and tightly packed neurofilament tracks^7^.

With an apparent molecular weight of 68 kDa, NFL is the smallest of the three core neurofilament subunits and is a nucleator of neurofilament assembly^8,9^. In common with the other neurofilament subunits, NFL is composed of three distinct domains. A short globular head domain at the N terminus of NFL is followed by a central α-helical rod domain and a C-terminal tail domain^10-12^. The N-terminal head domain contains sites at which posttranslational modifications involved in the regulation of neurofilament assembly occur and the rod domain contains α-helices responsible for the lateral interactions between neurofilament subunits and neurofilament assembly; the short, disordered C-terminal tail domain is also involved in the regulation of neurofilament assembly^7,13,14^. The roles of NFL overlap with the roles of neurofilaments in general. They provide mechanical support to axons, regulate axonal radial growth and caliber, and modulate axonal conduction velocity^1,7^. Apart from structural roles, an increasing number of studies suggest that neurofilaments have a role in the regulation of synaptic function and plasticity^15-20^. On its own, NFL is necessary for the assembly of neurofilaments^8,9^ and their interaction with other cytoskeletal elements^2,21,22^, myosin Va^23^, and *N*-methyl-D-aspartate receptors^15,19,24,25^. Furthermore, NFL is implicated in the pathophysiology of various diseases, indicating its importance for neuronal health. Several NFL mutations have been identified as a cause of Charcot–Marie–Tooth peripheral neuropathy^26-30^, and disruption to NFL metabolism and trafficking is associated with the pathophysiology of other diseases such as amyotrophic lateral sclerosis (ALS)^1^ and giant axon neuropathy^31^. Moreover, abnormal interactions between NFL and synaptic proteins have been linked with the synaptic dysfunction observed in some neuropsychiatric diseases^18,19^. Furthermore, the NFL released from injured neurons into extracellular fluids serves as a biomarker of neuronal damage^32-34^. Recent studies have shown that the amount of NFL in cerebrospinal fluid (CSF) and blood correlates with the severity of numerous diseases and conditions of the nervous system, including Alzheimer’s and Parkinson’s diseases, multiple sclerosis, ALS, and traumatic brain injury^33,35^.

Although these diseases differ in terms of their cause, pathophysiology, and clinical manifestation, a common factor contributing to their development and progression is oxidative stress-induced neuronal injury^36,37^. The main drivers of oxidative injury are reactive oxygen and nitrogen species such as superoxide anion, hydrogen peroxide, hydroxyl radical, peroxynitrite ion, and nitronium ion. Oxidative species cause neuronal injury by either direct oxidation of cellular proteins, lipids, and DNA, or through mitochondrial disruption and failure of energy production^37^. Energy deficiency causes Ca^2+^ ions to accumulate in the cytoplasm, activating Ca^2+^-dependent proteases such as calpains, which further contribute to neurodegeneration by degrading various proteins in the cell^38^. In addition, reactive nitrogen species cause damage to the cell through protein nitration, which disrupts protein structure and function^39^. During the oxidative injury of neurons, neurofilaments are affected in several ways. On the one hand, they are subjected to modifications such as changes in phosphorylation patterns, oxidation, nitration^1^, and *S*-nitrosylation^40^. These modifications disrupt the secondary structure of neurofilaments, their interactions, assembly, and axonal transport, leading to their aggregation and perikaryal accumulation. On the other hand, activated calpain proteases induce the proteolytic degradation of neurofilaments^41-43^. Products of this cleavage are potentially the ones released from the cell, contributing to the NFL levels detected in the CSF and blood of patients with neurological diseases^33^.

Despite its importance for neuronal homeostasis and its association with various neurological diseases, little is known about NFL beyond its classical role in structural support. Several studies have shown that NFL interacts with synaptic receptors, suggesting that it could also be involved in the regulation of neuronal function and activity^15,19,24^, but these roles are yet to be thoroughly investigated. Furthermore, although it is proposed as a biomarker of neuronal injury and a predictor of disease progression, the exact mechanism and functional implications of NFL release into the CSF and blood are not known^32-34^. In addition, although it is known that NFL is oxidized, nitrated, and degraded during oxidative injury, the downstream effects of these events on neurons remain elusive. To gain more information about these events, we investigated the response of NFL to oxidative injury induced in mouse cortical neurons *in vitro*. We found that during oxidative injury *in vitro*, NFL localizes in the nucleus of neurons. This accumulation is dependent on the activity of calpain proteases, as treatment with calpain inhibitors reduces the NFL signal in the nucleus. In addition, our results suggest that the NFL tail domain, or its fragment, localizes in the nucleus after being catalytically cleaved from the rest of the NFL. Finally, we demonstrate that after entering the nucleus the tail domain of NFL interacts with the DNA of living cells.

## Results

### Neurofilament light chain accumulates in neuronal nuclei upon nitric oxide-induced injury

To investigate the response of NFL to oxidative stress-induced injury, we treated primary mouse cortical neurons with a nitric oxide donor, spermine NONOate, or with the control compound sulfo NONOate, which does not release nitric oxide. After acute treatment with sulfo NONOate or spermine NONOate, neurons were fixed and stained with anti-NFL antibody. Anti-NFL antibody staining of neurons treated with the control compound showed the expected neurofilament morphology, that is, a strong fluorescence intensity in axons and lower intensity in cell bodies and dendrites (**Fig. 1a**). On the other hand, in neurons that were treated with the nitric oxide donor, anti-NFL antibody staining revealed NFL accumulation in what appeared to be the nuclei of the injured neurons (**Fig. 1b**). Co-staining with the nuclear dye SYTO16 (**Fig. 1a,b**) confirmed that this accumulation of NFL occurred in the nuclei of injured neurons, as the anti-NFL signal co-localized with that of SYTO16 (**Fig. 1b**). This is further exemplified by fluorescence intensity line profiles, which showed no increase of the NFL signal in the nuclei of control neurons, and a strong increase in the nuclei of neurons treated with the nitric oxide donor (**Fig. 1a,b**).

**Fig. 1.**
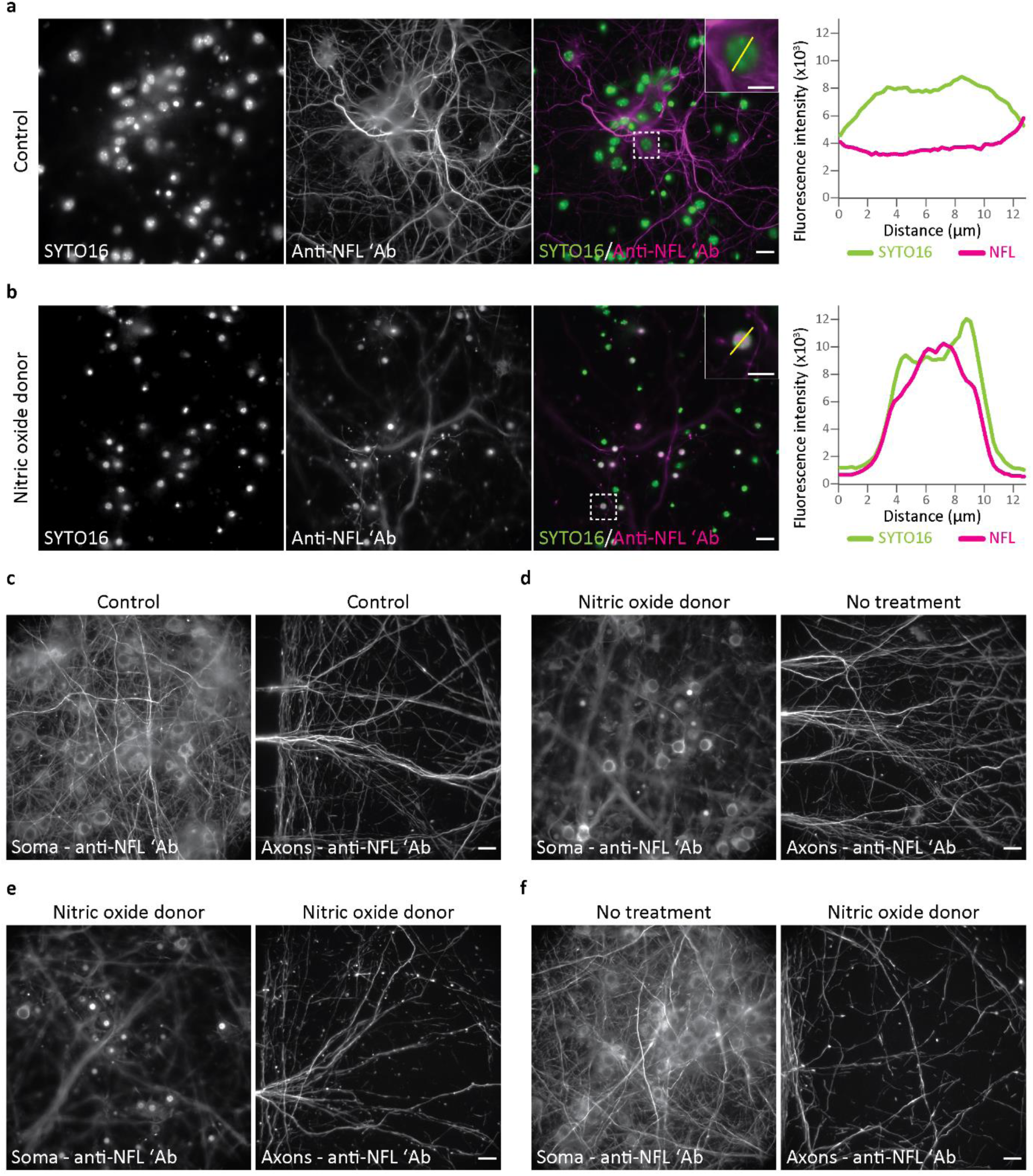
Localization of neurofilament light chain (NFL) in mouse cortical neurons (MCNs) after treatment with a nitric oxide donor. **a**,**b:** MCNs were treated at day *in vitro* (DIV) 12 for 3 h with either 500 μM sulfo NONOate (control; **a**) or 500 μM spermine NONOate (nitric oxide donor; **b**). After treatment, neurons were fixed, and stained with anti-NFL antibody and nuclear dye SYTO16. Graphs on the right show line profile fluorescence intensity measurements for both SYTO16 and anti-NFL signals. Fluorescence intensities are plotted as absolute gray values of raw 16-bit depth images. The regions of interest outlined by dashed boxes contain the nuclei used for the line profile measurement, expanded views of which are shown inset. **c−f:** MCNs were grown in compartmentalized microfluidic devices to separate axons from neuronal bodies and dendrites. At DIV 10, both bodies and axons were treated with 1 mM sulfo NONOate (control; **c**), or only bodies (**d**), both bodies and axons (**e**), or only axons (**f**) were treated with 1 mM spermine NONOate (nitric oxide donor). After 3 h of treatment, neurons were fixed and then stained with anti-NFL antibody. Images were acquired on a widefield microscope. Scale bars: 20 μm (**a−f**), 10 μm (images inset in **a** and **b**).

To investigate a potential dose dependency of the observed accumulation of NFL, we then treated neurons with spermine NONOate at various concentrations. These experiments showed that nuclear accumulation of NFL was induced with a spermine NONOate concentration as low as 25 μM and that the percentage of NFL-positive nuclei increased as the concentration of the nitric oxide donor was increased, reaching a 60% plateau at concentrations of 150 μM and greater (**Supplementary Fig. 1a,b**). Moreover, a time-course study revealed that NFL accumulates in the nuclei as early as 30 min after treatment with 250 μM spermine NONOate began (**Supplementary Fig. 1c**).

To exclude the possibility of nonspecific antibody binding, we performed immunocytochemical labeling of injured neurons with three other anti-NFL antibodies. In addition to the anti-NFL antibody used in the initial experiments, which targets the NFL tail domain (clone DA2; Merck, cat. no. MAB1615), we stained control and injured neurons with antibodies targeting NFL head (**Supplementary Fig. 2a,b**), NFL rod and tail (**Supplementary Fig. 2c,d**), or NFL tail (**Supplementary Fig. 2e,f**) domains. The results showed that the antibody that binds an epitope in the NFL head domain does not detect accumulated nuclear NFL (**Supplementary Fig. 2b**), whereas the antibody binding an unspecified epitope in the rod and tail regions of NFL does, as evidenced by a clear signal in the nucleus (**Supplementary Fig. 2d**). Surprisingly, the last of the three tested antibodies did not stain accumulated NFL in the nucleus (**Supplementary Fig. 2f**), even though its epitope should overlap with the epitope of the anti-NFL clone DA2 antibody, which reproducibly detects nuclear NFL accumulation (**Fig. 1, Supplementary Fig. 1**). Although inconsistent in terms of which NFL domain accumulates in the nucleus, these results show that nuclear NFL accumulation can be detected with at least one other anti-NFL antibody (**Supplementary Fig. 2d**). Again, this was confirmed by SYTO16 co-staining and line profile fluorescence intensity measurements (**Supplementary Fig. 2**).

Further experiments revealed that nuclear accumulation of NFL is not specific to mouse cortical neurons, as it was also detected in mouse hippocampal, rat cortical, and human induced pluripotent stem cell (IPSC)-derived neurons (**Supplementary Fig. 3**). In addition to NFL, we tested whether other neurofilament subunits accumulate in the nucleus upon nitric oxide treatment. Post-injury immunocytochemical labeling revealed that neither NFM nor nonphosphorylated NFH accumulate in the nuclei of injured neurons (**Supplementary Fig. 4**).

In a further investigation of these findings, we wondered whether NFL accumulates in the nucleus after injury of different neuronal compartments. For this purpose, we cultured mouse cortical neurons in microfluidic devices that allow spatial separation as well as independent injury of somato-dendritic and axonal compartments. Consistent with the results from non-compartmentalized neuronal cultures, treatment of neurons cultured in microfluidic devices with the control compound caused no disruption to neurofilament networks and localization (**Fig. 1c**). Furthermore, injury of somato-dendritic and axonal compartments alone, or in combination, showed that NFL accumulated in the nuclei only if neuronal bodies were injured (**Fig. 1d,e**). Acute injury to only axons had no effect on the NFL in neuronal bodies and no accumulation of NFL in the nucleus was observed (**Fig. 1f**).

### Neurons with nuclear NFL accumulation undergo necrosis

After the initial experiments that described the nuclear accumulation of NFL upon neuronal injury, we studied whether neurons that accumulate NFL in the nucleus are in the process of dying. After we induced acute injury of primary mouse cortical neurons with nitric oxide, we performed a terminal deoxynucleotidyl-transferase dUTP nick end labeling (TUNEL) assay^44,45^ to investigate if the neurons are in the late stages of apoptosis. The results showed that both control and injured neurons accumulating nuclear NFL were TUNEL-negative, suggesting that injured neurons were not undergoing late-stage apoptosis (**Fig. 2a,b**). Control experiments in which control and injured neurons were treated with deoxyribonuclease I (DNase I) yielded a TUNEL-positive signal, validating the results of the assay (**Supplementary Fig. 5a,b**).

**Fig. 2.**
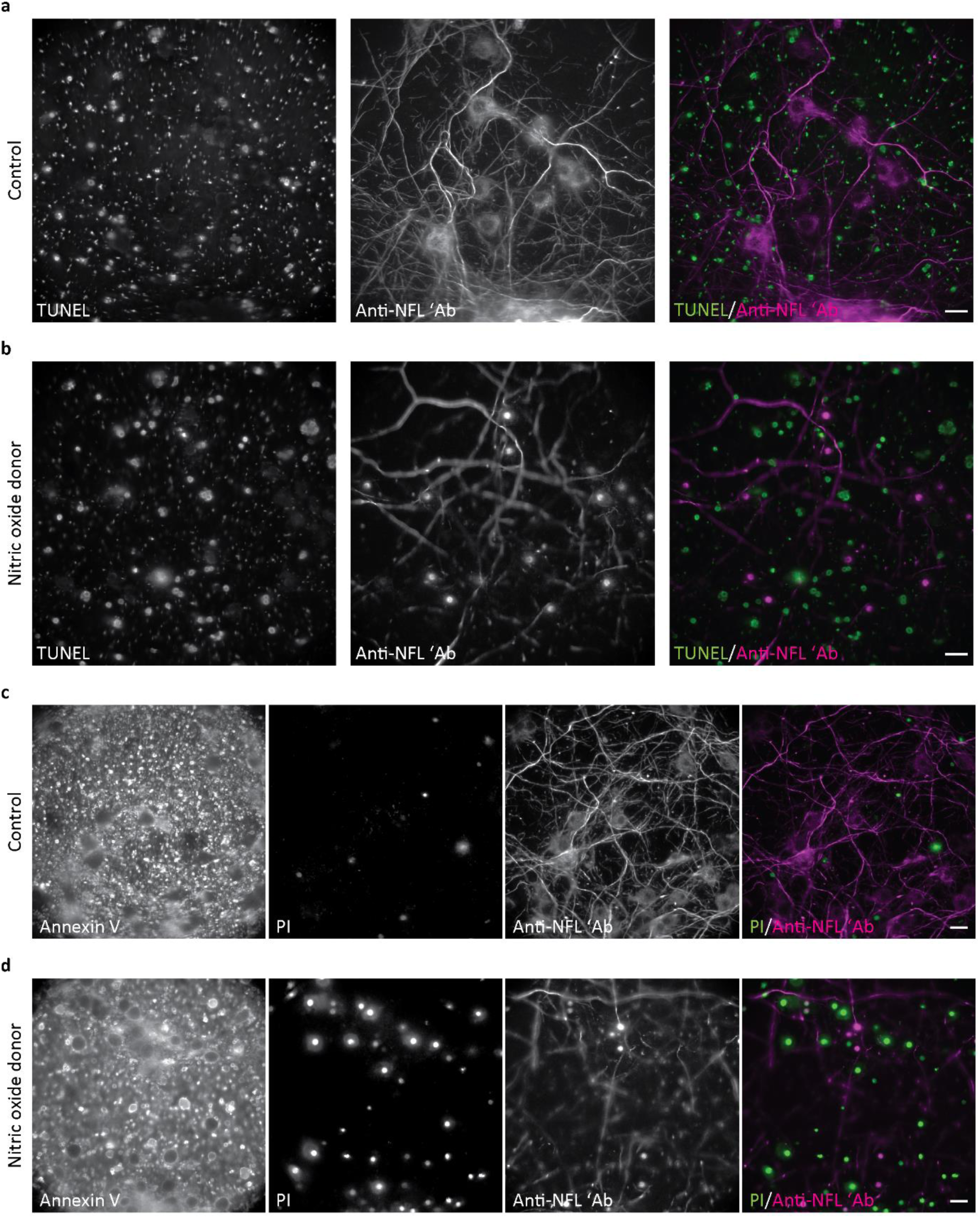
Detection of apoptotic or necrotic cell death in neurons treated with a nitric oxide donor. **a–d:** MCNs were treated at DIV 10 (**a**,**b**) or DIV 12 (**c**,**d**) with either sulfo NONOate (control; **a**,**c**) or spermine NONOate (nitric oxide donor; **b**,**d**). **a**,**b:** After treatment with 250 μM NONOates for 3 h, neurons were fixed and labeled in a terminal deoxynucleotidyl transferase dUTP nick end labeling (TUNEL) assay for the detection of late-stage apoptosis, and with anti-NFL antibody. **c**,**d:** After treatment with 500 μM NONOates for 2.5 h, neurons were labeled with annexin V and propidium iodide (PI) for the detection of early-stage apoptosis or necrosis, respectively, and imaged live. Then, neurons were fixed and stained with anti-NFL antibody. Images were acquired on a widefield microscope. Scale bars: 20 μm.

Next, we wanted to test if neurons with accumulated nuclear NFL are in the initial stages of apoptosis or undergoing necrosis. To this end, we again induced injury of primary mouse cortical neurons and performed an annexin V/propidium iodide (PI) assay. Annexin V specifically binds the phosphatidylserine that is exposed on the cell surface during the initial stages of apoptosis^46^. By contrast, the otherwise cell-impermeable PI enters the cell and binds its DNA in the nucleus only if the integrity of the cell membrane is compromised, as typically happens during necrosis^46^. We treated neurons with either the control compound sulfo NONOate or with the nitric oxide-releasing spermine NONOate, performed annexin V and PI staining, and imaged the live cells with widefield microscopy. As expected, control neurons showed only the background staining of cellular debris and no annexin V was detected on the cell membrane (**Fig. 2c**). Correspondingly, PI-positive staining was rare in control cultures, again, most signal derived from the staining of cellular debris (**Fig. 2c**). In contrast to control neurons, nitric oxide-treated neurons were positive for both annexin V and PI (**Fig. 2d**). Dual-positive annexin V/PI staining suggests that neurons are either in the late stages of apoptosis when rupturing of the cell membrane occurs, or undergoing necrosis. Considering that injured neurons are not in the late stage of apoptosis (**Fig. 2b**), and that phosphatidylserine is also exposed in the outer layer of the membrane during necrosis^46,47^, our results suggest that neurons treated with nitric oxide undergo necrosis. Anti-NFL staining after live-cell imaging and fixation confirmed that injured annexin V/PI-positive neurons accumulate NFL in the nucleus (**Fig. 2d**). Interestingly, neurons with nuclear NFL accumulation had a low PI fluorescence intensity, suggesting that NFL might be interfering with PI binding to the DNA. Additional control experiments in which neurons were treated with 10 μM staurosporine (a positive control for induction of apoptosis) or with 0.1% saponin (a positive control for induction of necrosis) confirmed the efficiency of the annexin V/PI assay (**Supplementary Fig. 5c**,**d**). Taken together, these results suggest that injured neurons accumulating NFL in the nucleus are in neither the initial nor the late stages of apoptosis, and undergo necrotic cell death.

### Nuclear accumulation of NFL is not specific to nitric oxide-induced injury and can be prevented by calpain activity inhibition

Since all of our previous experiments centered on injury induced by nitric oxide, we wanted to investigate if nuclear accumulation of NFL arises specifically from that. Thus, we also injured primary mouse cortical neurons with hydrogen peroxide, carbonyl cyanide 3-chlorophenylhydrazone (CCCP), and monosodium glutamate, and compared them to healthy neurons (**Fig. 3a**). Hydrogen peroxide is a reactive oxygen species that introduces oxidative damage to cellular structures^36^, whereas CCCP stimulates the accumulation of reactive oxygen species in the cell by lowering mitochondrial potential^48^. By contrast, monosodium glutamate induces excitotoxicity by increasing neuronal activity^49^. After injury with hydrogen peroxide and CCCP, we observed strong accumulation of NFL in neuronal nuclei (**Fig. 3b,c**) similar to that resulting from nitric oxide injury (**Fig. 1b**). Conversely, after monosodium glutamate treatment, accumulation of NFL in the nucleus was not as prominent, and could be observed only after adjusting image brightness and contrast to very low values (**Fig. 3d, arrows**). These results suggest that nuclear accumulation of NFL is not specific to nitric oxide-induced injury but rather a common outcome of oxidative damage to the cell.

**Fig. 3.**
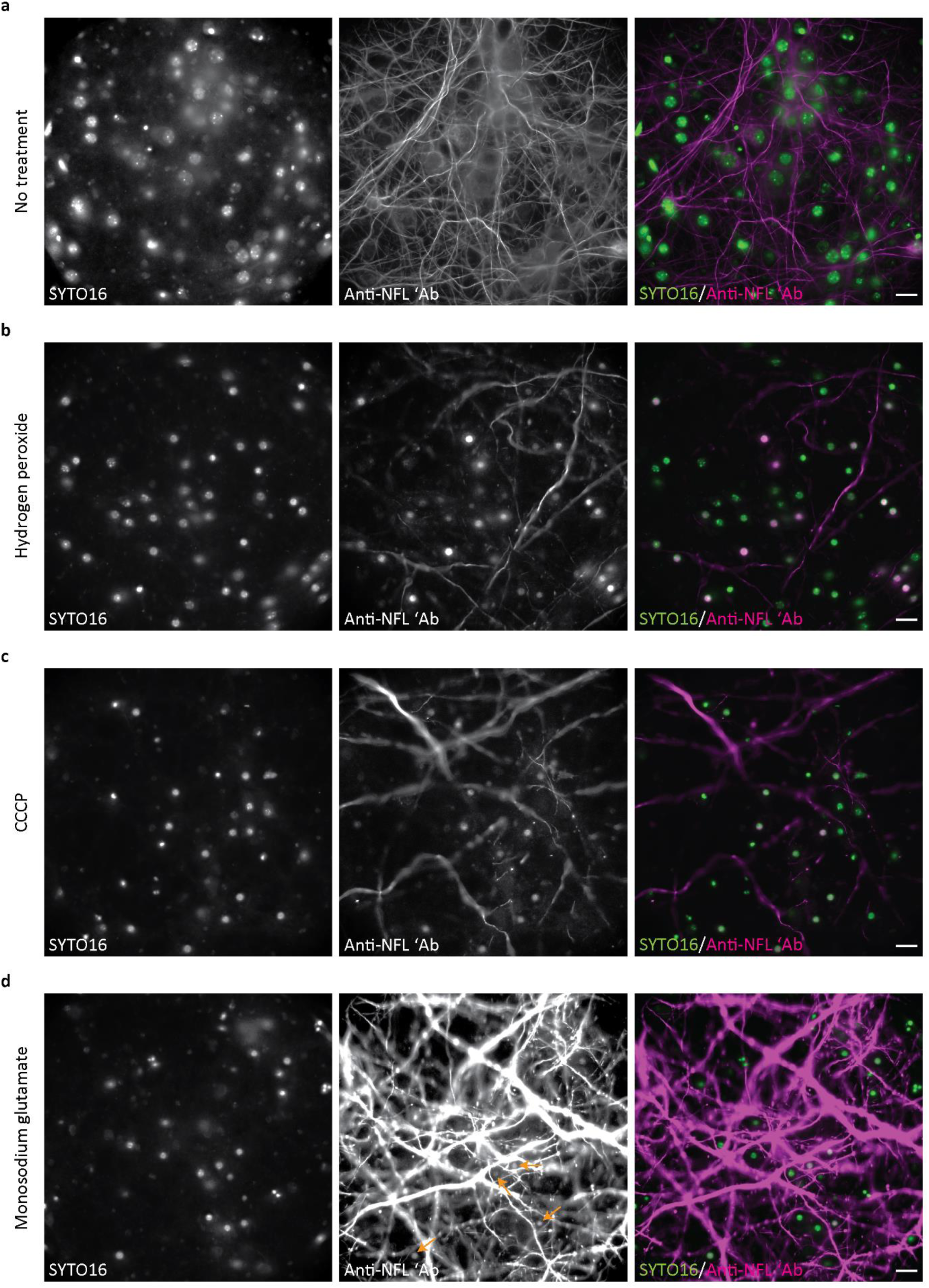
Localization of NFL in neurons after treatment with additional oxidative stress-inducing compounds. MCNs were either not treated (**a**) or were treated with 1 mM hydrogen peroxide for 3 h (**b**), 25 μM carbonyl cyanide 3-chlorophenylhydrazone (CCCP) for 3 h (**c**), or 100 μM monosodium glutamate for 6 h (**d**). Neurons were then fixed, and stained with anti-NFL antibody and the nuclear dye SYTO16. Images were acquired on a widefield microscope. Arrows in panel **d** point at the nuclei with accumulation of NFL. Scale bars: 20 μm.

We next wanted to investigate if accumulation of NFL in the nucleus depends on the activity of calpain proteases. It is well known that oxidative stress and injury cause the activation of calpain proteases, which degrade various proteins in the cell, including NFL^41-43^. Accordingly, we investigated the effect calpain inhibition has on the oxidative injury-induced accumulation of NFL in the nucleus. We pre-treated primary mouse cortical neurons with various calpain or caspase inhibitors and then induced injury with nitric oxide. We used calpain inhibitor III (MDL 28170), which is a selective and reversible inhibitor, and EST, an irreversible calpain inhibitor^50^. As a control, we used the pan-caspase inhibitor emricasan to inhibit the activity of caspases^51^ which should not have an effect on NFL^52^. Widefield imaging of nuclear NFL accumulation showed that pre-treatment with calpain inhibitors reduced the intensity of NFL staining in the nucleus of injured neurons, whereas pre-treatment with caspase inhibitor expectedly had no effect (**Fig. 4a**). To confirm these findings, we quantified the fluorescence intensity of antibody-labeled NFL in neuronal nuclei (**Fig. 4b**). However, as the nuclei of cultured neurons are situated above the level of dendrites and axons, some of the nuclei in the control sulfo-NONOate-treated group gave a false-positive signal for NFL arising from the out-of-focus neurofilaments (**Supplementary Fig. 6a,b**). In addition, since oxidative injury induces degradation of NFL, control neurons had a more complex neurofilament network with a higher anti-NFL fluorescence intensity than the injured neurons. Consequently, these bright out-of-focus neurofilaments contributed significantly to the high false-positive signal in the nuclei of some control cells. Furthermore, as only a maximum of 60% of neurons accumulate NFL upon nitric oxide-induced injury (**Supplementary Fig. 1b**) and due to the NFL degradation during injury, each of the nitric oxide-treated groups contained nuclei with a low NFL signal. To avoid the influence of these nuclei and out-of-focus background signals on the analysis, we determined a cutoff value that corresponded to the median value of the control group (**Fig. 4b, dashed line**). Quantification analysis of the data above the cutoff showed that treatment with nitric oxide significantly increased the intensity of NFL in the nucleus in comparison to non-injured control neurons. Furthermore, treatment of neurons with calpain inhibitors, or their combination with the caspase inhibitor, significantly reduced the intensity of nuclear NFL in comparison to neurons that were treated with the nitric oxide only (**Fig. 4b, Supplementary Table 1**), confirming the observations made by microscopy (**Fig. 4a**). In addition, because there was no significant difference between neurons treated with the caspase inhibitor and neurons treated only with the nitric oxide donor, quantitative analysis confirmed the finding by microscopy that pre-treatment with a caspase inhibitor had no significant effect on NFL accumulation in neuronal nuclei (**Fig. 4a,b, Supplementary Table 1**). Furthermore, there was no significant difference between treatments with calpain inhibitor III and EST either alone or in combination, nor between treatments with calpain inhibitors alone and in combination with the caspase inhibitor (**Fig. 4b, Supplementary Table 1**). These results show that both reversible and irreversible calpain inhibitors significantly reduce nuclear NFL accumulation, and that their combination shows neither additive nor synergistic effects on NFL accumulation in the nucleus. Moreover, these results support the finding that the caspase inhibitor emricasan does not have an impact on NFL accumulation in the nucleus resulting from nitric oxide-induced injury.

**Fig. 4.**
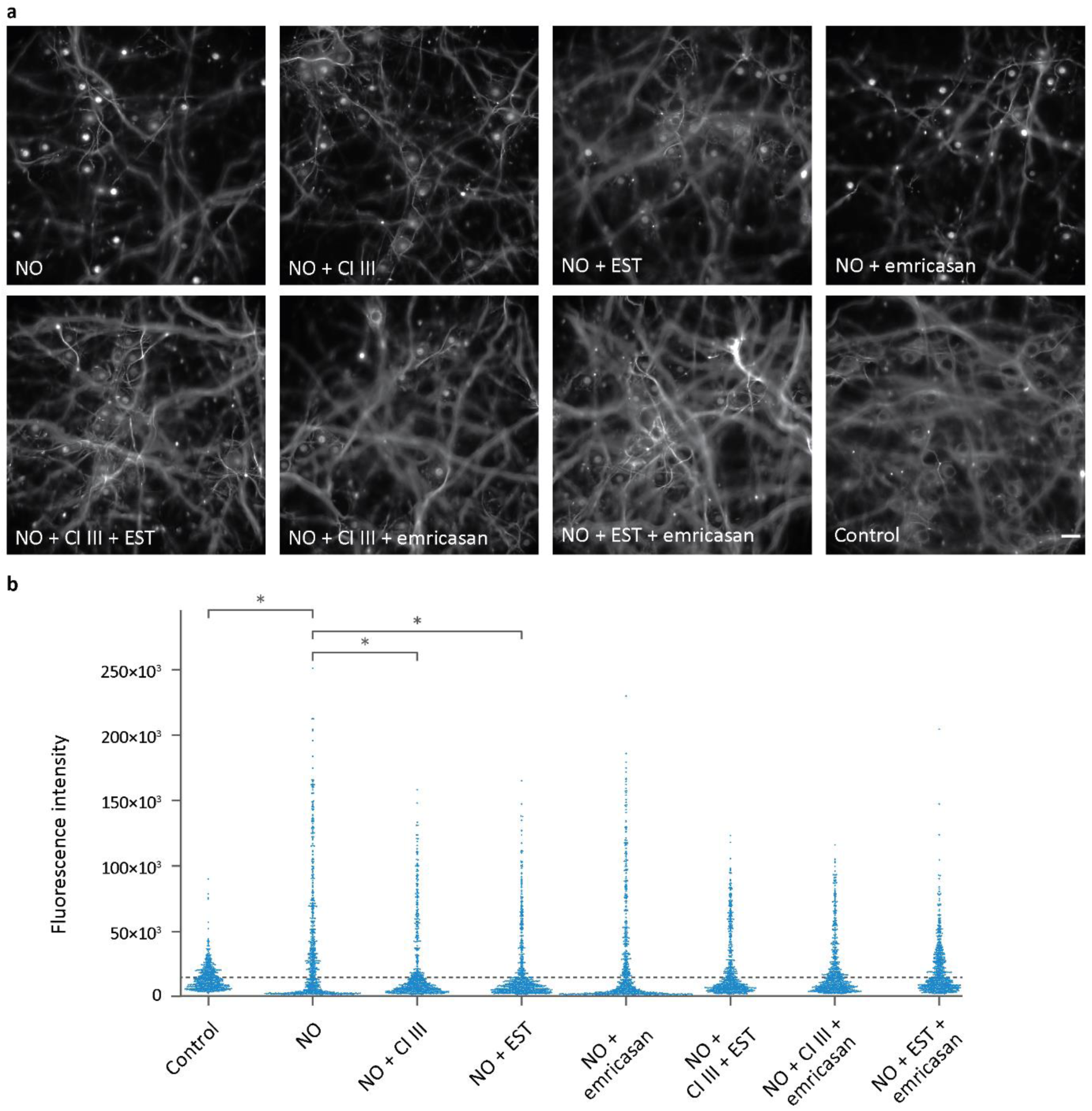
Quantification of nuclear localization of NFL after pre-treatment with calpain or caspase inhibitors and treatment with nitric oxide. **a:** DIV-12 MCNs were pre-treated for 1 h with 100 μM calpain inhibitor III (CI III), 100 μM calpain inhibitor EST or 20 μM caspase inhibitor (emricasan), then treated for 3 h with either 500 μM sulfo NONOate (control) or 500 μM spermine NONOate (NO). Neurons were then fixed, stained with anti-NFL antibody and the nuclear dye SYTO16, and imaged with widefield microscopy. Scale bar: 20 μm. **b**: Quantification of nuclear NFL fluorescence intensity, performed on the data set that was acquired as described for panel **a**. Nuclear NFL fluorescent labeling intensity was measured in Fiji/ImageJ software and used to determine the corrected total cell fluorescence (CTCF) intensity values. Absolute values of CTCF are plotted on the y axis. As not all the nuclei of injured neurons contained accumulated NFL, each of the nitric-oxide-treated groups had nuclei with a low NFL signal. For this reason, only the nuclei above a threshold corresponding to the median of the control group (dashed line) were analyzed in each group. CTCF values were compared by running a Kruskal–Wallis non-parametric test, followed by a Dunn–Bonferroni post-hoc correction for multiple comparisons. Data were collected from three independent experiments, with 10 images acquired per group per experiment. A total of 2342 nuclei were analyzed; the number of nuclei per group was as follows: control *n*=272; nitric oxide *n*=361; nitric oxide + calpain inhibitor III *n*=246; nitric oxide + EST *n*=251; nitric oxide + emricasan *n*=270; nitric oxide + calpain inhibitor III + EST *n*=256; nitric oxide + calpain inhibitor III + emricasan *n*=323; nitric oxide + EST + emricasan *n*=363. The most relevant significant differences are indicated with an asterisk (*). Details of the statistical analysis and all significant differences between groups are given in Supplementary Table 1.

### Live- and fixed-cell imaging of exogenously expressed NFL during oxidative neuronal injury

As our previous experiments relied on determining the localization of endogenous NFL after injury, fixation, and immunocytochemical labeling, they were not suitable for studying the dynamics of NFL translocation in live cells as injury occurs. To that end, we transfected neurons with C-terminally tagged NFL (NFL-GFP) and monitored its localization in cells exposed to a nitric oxide donor. Live-cell imaging of neurons expressing NFL-GFP showed no change in the neurofilament network upon treatment with the control compound (**Fig. 5a**). However, in cells exposed to nitric oxide, we observed that GFP is cleaved from NFL, distributes uniformly throughout the cytoplasm, and is no longer a marker of NFL localization in injured neurons (**Fig. 5b, Supplementary Video 1**).

To overcome this setback, we first examined if the C-terminal region of NFL contains putative calpain cleavage sites that might be responsible for the separation of NFL and GFP observed during live-cell imaging. Using the two previously published algorithms for calpain cleavage site prediction (DeepCalpain^53^ and GPS-CCD^54^), we identified several putative calpain cleavage sites in the C-terminal region of NFL. We focused on the sites localized between the epitope recognized by the anti-NFL antibody clone DA2 (amino acids A441–A461) and the C terminus of NFL (**Supplementary Table 2**). In addition, we identified several putative cleavage sites in the GFP sequence that could also be responsible for the observed NFL–GFP separation. In an attempt to render our NFL-GFP construct calpain-insensitive and prevent the cleavage taking place, we removed the C-terminal sequence of NFL after the residue A461. NFL(ΔA461–D543)-GFP expressed well in neurons, incorporated successfully into the neurofilament network, and was indistinguishable from the full-length NFL-GFP (**Fig. 5c,d**). During the exposure to the control compound sulfo NONOate, there was no change in the neurofilament network that contained NFL(ΔA461–D543)-GFP, similar to the full-length NFL-GFP (**Fig. 5c**). Conversely, during the treatment with nitric oxide, NFL(ΔA461–D543)-GFP was not cleaved by calpain proteases to the same extent as the full-length NFL-GFP. Consequently, we could observe the increase of GFP fluorescence intensity in the nucleus during the course of the treatment (**Fig. 5d**). However, in many neurons that were monitored throughout the experiment, NFL(ΔA461–D543) and GFP still separated, and nuclear accumulation was untraceable. These results suggest that calpain cleavage of NFL(ΔA461–D543)-GFP still occurs despite the deletion of putative cleavage sites in the NFL sequence. Previously identified sites at the beginning of the GFP sequence might be responsible for this cleavage. To further eliminate the possibility of cleavage in the tag itself, we added a C-terminal FLAG tag to both NFL and NFL(ΔA461–D543). Despite its incompatibility with live imaging, we selected the FLAG tag since it contained no putative calpain cleavage sites. We expressed NFL-FLAG and NFL(ΔA461–D543)-FLAG constructs in primary mouse cortical neurons and treated them with either sulfo NONOate or spermine NONOate. Cells were then fixed and stained with anti-NFL and anti-FLAG antibodies and also with the nuclear dye SYTO16. As in previous experiments, control-treated neurons expressing either NFL-FLAG or NFL(ΔA461–D543)-FLAG had a normal neurofilament network (**Fig. 6a,c**). After treatment with nitric oxide, only six out of 51 transfected neurons expressing the full-length NFL-FLAG contained a FLAG-positive nucleus (**Fig. 6b**). By contrast, almost all injured neurons expressing NFL(ΔA461–D543)-FLAG, had a clear FLAG signal in the nucleus (42 out of 51 transfected cells; **Fig. 6d**). This was further confirmed by the fluorescence intensity line profiles of anti-FLAG, anti-NFL, and SYTO16 staining in the nucleus of control and injured neurons (**Fig. 6**). These results demonstrate that exogenously expressed recombinant NFL behaves as endogenous NFL during injury, and suggest that they are both subject to at least one calpain cleavage in the C-terminal tail domain.

**Fig. 5.**
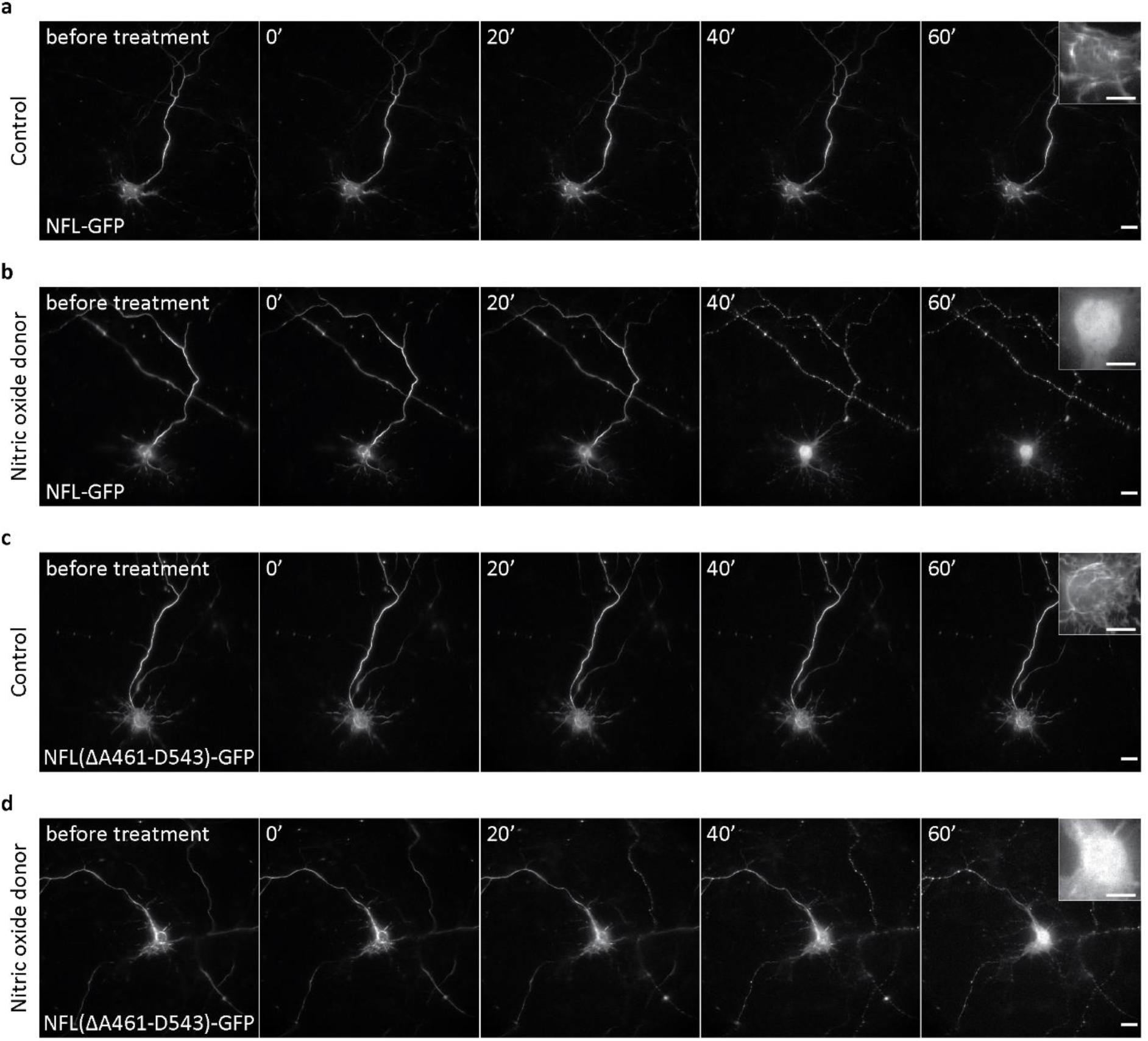
Live imaging of NFL translocation to the nucleus during nitric oxide-induced injury in neurons. **a–d:** MCNs were transfected at DIV 8 with constructs bearing either NFL-GFP (**a**,**b**) or NFL(ΔA461–D543)-GFP (**c**,**d**). After 3 days of expression, neurons were treated with 500 μM of either control compound sulfo NONOate (**a**,**c**) or nitric oxide donor spermine NONOate (**b**,**d**) and imaged live during injury with widefield microscopy. Scale bars: 20 μm (**a−d**), 10 μm (images inset in **a−d**).

**Fig. 6.**
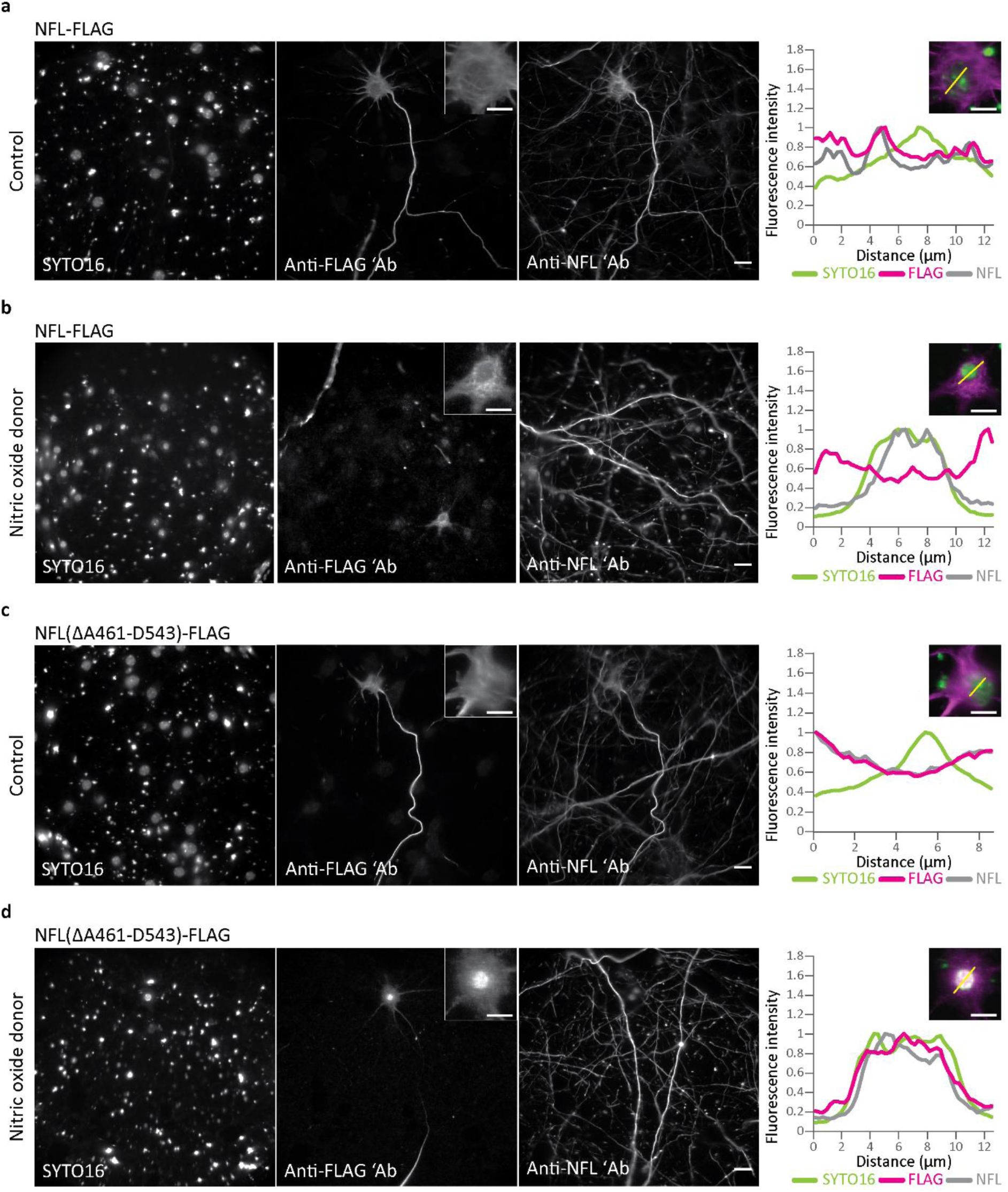
Localization of NFL-FLAG and NFL(ΔA461–D543)-FLAG during nitric oxide-induced injury in neurons. **a–d:** MCNs were transfected at DIV 8 with constructs bearing either NFL-FLAG (**a**,**b**) or NFL(ΔA461–D543)-FLAG (**c**,**d**). After 3 days of expression, neurons were treated for 3 h with either 500 μM sulfo NONOate (control; **a**,**c**) or 500 μM spermine NONOate (nitric oxide donor; **b**,**d**). Neurons were then fixed, stained with anti-FLAG and anti-NFL antibodies and the nuclear dye SYTO16, and imaged with widefield microscopy. Graphs on the right show line profile fluorescence intensity measurements for SYTO16, anti-FLAG, and anti-NFL signals. Fluorescence intensities were normalized to the highest intensity value of each channel. Merged images of SYTO16 (green) and anti-FLAG (magenta) stainings (shown inset above the graphs) show the line regions of interest from which the respective fluorescence intensities were measured. Scale bars: 20 μm (**a−d**), 10 μm (images inset in **a−d**).

### The C-terminal tail domain of NFL accumulates in the nuclei of healthy neurons

Having observed that calpain inhibition prevented NFL accumulation in the nucleus (**Fig. 4**) and that removing putative calpain cleavage sites from the NFL tail domain prevented the separation of NFL from its FLAG tag (**Fig. 6**), we hypothesized that a proteolytic fragment of NFL and not the full-length NFL accumulates in the nucleus upon injury. Based on our initial experiments utilizing the anti-NFL antibody that recognizes an epitope in the tail domain of NFL (**Fig. 1**), and based on the labeling with other antibodies that recognize different NFL domains (**Supplementary Fig. 2**), we suspected that NFL rod or tail domains, or their proteolytic fragments, accumulate in the nucleus during injury. To test this hypothesis, we transfected healthy neurons with constructs encoding full-length NFL or NFL head, rod, or tail domains. All of these domains were tagged with a C-terminal FLAG, which we used to detect the NFL domains in transfected cells (**Fig. 7a**). The results showed that full-length NFL-FLAG was expressed well and interacted with other neurofilament subunits to form neurofilaments in healthy neurons (**Fig. 7b**). In addition, the transfection efficiency for introducing these NFL domains appeared lower than that for the full-length NFL. In contrast to the full-length NFL, none of the NFL domains were incorporated into the neurofilament network; all disrupted neurofilament assembly and were distributed uniformly in the neuronal cytoplasm, with NFL head and tail domains also observed in the nucleus (**Fig. 7c−e**). Although both NFL head and tail domains localized in the nuclei of healthy neurons, NFL tail domain showed a tendency to accumulate and prevent nuclear labeling with the SYTO16 dye (**Fig. 7e**). This was further illustrated by the line profile fluorescent intensity measurement, which showed a lower SYTO16 signal at the location of NFL tail accumulation in the nucleus (**Fig. 7e**). In contrast to the tail, NFL head localization in the nucleus did not affect SYTO16 labeling (**Fig. 7c**).

**Fig. 7.**
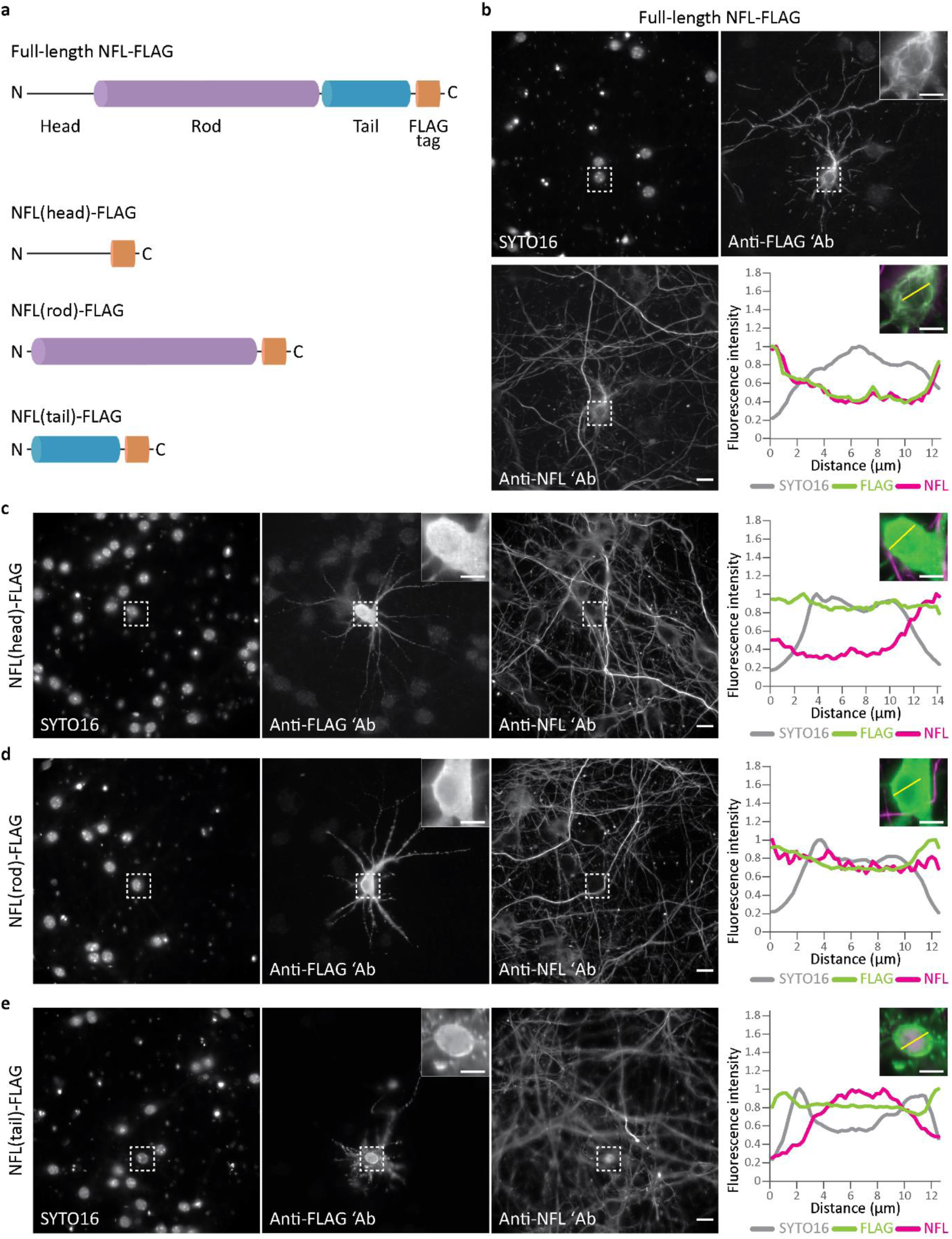
Localization of recombinant full-length NFL and NFL domains in healthy neurons. **a:** Schematic representation of the constructs used in this experiment. Full-length NFL contains all NFL domains and a FLAG tag at the C terminus. Other constructs contain only head, rod, or tail domains of NFL tagged with a C-terminal FLAG tag. **b−e:** MCNs were transfected at DIV 8 with full-length NFL-FLAG (**b**), NFL(head)-FLAG (**c**), NFL(rod)-FLAG (**d**), or NFL(tail)-FLAG (**e**). After 3 days of expression, neurons were fixed, stained with anti-FLAG and anti-NFL antibodies and the nuclear dye SYTO16, and imaged with widefield microscopy. Graphs on the right show line profile fluorescence intensity measurements for SYTO16, anti-FLAG, and anti-NFL signals. Fluorescence intensities were normalized to the highest intensity value of each channel. Merged images of anti-FLAG (green) and anti-NFL (magenta) stainings (shown inset above the graphs) show the line regions of interest from which the respective fluorescence intensities were measured. Expanded views of regions of interest (dashed boxes) are shown inset in anti-FLAG panels. Scale bars: 20 μm (**b−e**), 10 μm (images inset in **b−e**).

Since overexpression of neurofilament subunits could unbalance the ratio of neurofilament subunits in a cell, resulting in disruption of neurofilament assembly, we were concerned that overexpression of NFL domains would have the same effect and be the cause of the observed accumulation. We thus performed an additional set of experiments in which either full-length NFL or NFL domains were co-transfected with NFM (**Supplementary Fig. 7**). The results showed that co-expression of NFM could not rescue the disrupted neurofilament assembly caused by the recombinant NFL domains (**Supplementary Fig. 7b−d**). In addition, NFM co-expression did not prevent the NFL head and tail domains from accumulating in the nucleus, although fewer neurons exhibited nuclear accumulation of the NFL tail (**Supplementary Fig. 7c,d**).

In addition to experiments in primary neurons, we further investigated the localization and potential accumulation of NFL domains in the nuclei of ND7/23 mouse neuroblastoma and a rat neuron hybrid cell line. Due to their neuronal properties^55^, ND7/23 cells are frequently used as a model of primary neurons. We transfected ND7/23 cells with the full-length NFL and NFL domains with or without NFM (**Supplementary Fig. 8 and 9**). Results of these experiments were similar to those obtained with primary neurons in our earlier analogous experiments. After overexpression, NFL head and rod domains were distributed homogenously in the cytoplasm of ND7/23 cells, whereas the NFL tail domain accumulated both in the cytoplasm and in the nucleus, and prevented SYTO16 labeling (**Supplementary Fig. 8b−d**). However, in contrast to the results obtained in neurons, the NFL head domain accumulated primarily in the cytoplasm, exhibiting a low signal in the nuclei of ND7/23 cells (**Supplementary Fig. 8b**). The results of co-expressing NFL domains with NFM were the same as in primary neurons, except in the case of the NFL head domain that formed cytoplasmic aggregates after co-expression with NFM (**Supplementary Fig. 9**).

**Fig. 8.**
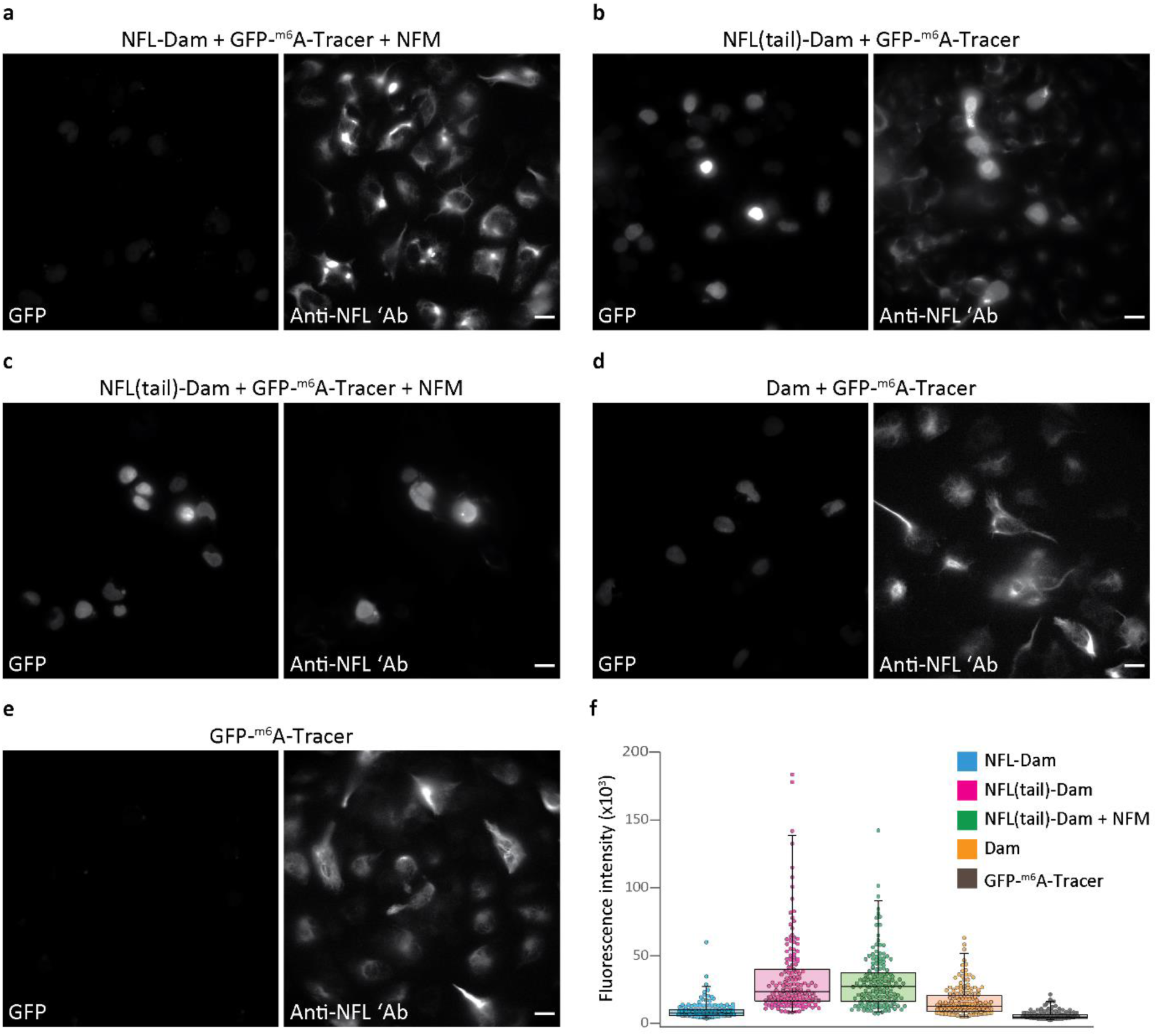
Tail domain of NFL interacts with the DNA in the nuclei of neuroblastoma ND7/23 cells. **a−e:** ND7/23 cells were transfected with constructs expressing the full-length NFL-Dam, NFM, and GFP-^m6^A-Tracer (**a**), or NFL(tail)-Dam and GFP-^m6^A-Tracer without (**b**) or with (**c**) NFM. Control cells were transfected with constructs expressing Dam and GFP-^m6^A-Tracer (**d**) or GFP-^m6^A-Tracer only (**e**). After incubation overnight, cells were fixed and stained with anti-NFL antibody. Images were acquired on a widefield microscope. The brightness and contrast of GFP images were scaled linearly and in the same way in each panel so that the differences in GFP fluorescence can be readily observed. **f:** Quantification of nuclear GFP fluorescence intensity, performed on the data set that was acquired as described for panels **a−e**. Nuclear GFP fluorescent labeling intensity was measured in Fiji/ImageJ software and used to determine the corrected total cell fluorescence (CTCF) intensity values. Absolute values of CTCF are plotted on the y axis. CTCF values were compared by running a Kruskal–Wallis non-parametric test, followed by a Dunn–Bonferroni post-hoc correction for multiple comparisons. Data were collected from two independent experiments, with 10 images acquired per group per experiment. A total of 941 nuclei were analyzed; the number of nuclei per group was as follows: NFL-Dam *n*=215; NFL(tail)-Dam *n*=170; NFL(tail)-Dam + NFM *n*=169; Dam *n*=221; GFP-^m6^A-Tracer *n*=166. Details of the statistical analysis are given in Supplementary Table 4. Scale bars: 20 μm.

### The NFL tail domain interacts with the DNA in the nuclei of neuroblastoma ND7/23 cells

After observing that endogenous NFL accumulated in the nuclei of injured neurons and that the recombinant tail domain of NFL accumulated in the nuclei of both primary neurons and ND7/23 cells, we wanted to investigate the biological significance of this accumulation. Since NFL accumulation in the nuclei of primary neurons decreased PI staining (**Fig. 2d**), and NFL tail accumulations prevented SYTO16 labeling (**Fig. 7e, Supplementary Fig. 8d**), we hypothesized that NFL tail binds the nuclear DNA after translocating into the nucleus during injury. To investigate whether NFL interacts with the genomic DNA in cells, we utilized “Dam identification” (DamID), a previously described method for studying protein–DNA interactions based on the DNA adenine methyltransferase (Dam) from *Escherichia coli*^56^. DamID involves making a fusion of the protein of interest with the Dam, which then methylates the N^6^ position of adenine within a GATC sequence if the protein of interest comes into close proximity with the DNA. The interaction of the protein of interest with the DNA is subsequently identified through the detection of DNA methylation. A microscopy-compatible approach for the detection of DNA methylation relies on the expression of a fusion between GFP and a DNA-binding fragment of the restriction enzyme DpnI (GFP-^m6^A-Tracer)^57^.

To test if NFL or its tail domain interact with DNA in the nuclei of neurons and ND7/23 cells, we created NFL-Dam and NFL(tail)-Dam C-terminal protein fusions. We transfected primary neurons with these constructs and with the GFP-^m6^A-Tracer, then fixed and stained with anti-NFL antibody. The results showed that the NFL(tail)-Dam underwent the expected nuclear accumulation and co-localized with the GFP signal in the nucleus (**Supplementary Fig. 10a**). However, the full-length NFL-Dam did not incorporate well into the neurofilament network in primary neurons and forms large cytoplasmic aggregates, even when co-expressed with NFM (**Supplementary Fig. 10b**). These results suggest that the addition of the Dam enzyme to the C terminus of NFL interferes with neurofilament assembly, thus rendering NFL-Dam fusion suboptimal for experiments in primary neurons.

We therefore focused our experiments on ND7/23 cells. We transfected ND7/23 cells with NFL-Dam or NFL(tail)-Dam constructs, together with the GFP-^m6^A-Tracer. In addition, we transfected control cells with constructs expressing Dam and GFP-^m6^A-Tracer, or only with the construct expressing GFP-^m6^A-Tracer. The results of these experiments showed that in contrast to neurons, in ND7/23 cells co-expressing NFM, the full-length NFL-Dam fusion integrated well into the neurofilament network (**Fig. 8a**). Moreover, ND7/23 cells that expressed NFL-Dam had a weak GFP signal in the nucleus, suggesting that the full-length NFL does not interact with the DNA (**Fig. 8a**). On the contrary, ND7/23 cells that expressed NFL(tail)-Dam had a strong GFP signal in the nucleus, indicating a high degree of DNA methylation (**Fig. 8b,c**). Furthermore, a strong GFP signal was present regardless of NFM co-expression, which co-localized with the accumulated NFL tail in the nucleus as detected by anti-NFL staining (**Fig. 8b,c**). Control cells that expressed Dam showed a low GFP signal in the nucleus, albeit higher than the cells that expressed NFL-Dam (**Fig. 8d**). As expected, cells that expressed only GFP-^m6^A-Tracer had an almost undetectable GFP signal in the nucleus (**Fig. 8e**). Quantification analysis (**Fig. 8f**) showed that the GFP fluorescence intensity was significantly higher in the nuclei of ND7/23 cells expressing NFL(tail)-Dam, either with or without NFM, compared to that in cells expressing full-length NFL-Dam or Dam alone. These results suggest that the NFL tail domain interacts with the genomic DNA after translocating into the nucleus of ND7/23 cells.

## Discussion

Here we report novel aspects of the response of NFL to the *in vitro* oxidative stress of neurons. First, we discovered that in response to nitric oxide-induced injury, NFL localizes in the nuclei of primary neurons. Previous studies on the effects of oxidative stress on neurofilaments have principally relied on assays with purified NFL^58-62^ or on tissues derived from patients with neurological disease^63^. These studies found that oxidative stress causes NFL aggregation^58-60,62,63^ and increased neurofilament susceptibility to calpain degradation^61^. In addition, several studies have shown oxidative stress-induced NFH hyperphosphorylation and subsequent neurofilament aggregation in cultured neurons^64,65^. However, in contrast to our results, these *in vitro* studies reported only perikaryal aggregation of neurofilaments caused by oxidative stress and not the detection of NFL in the nuclei of affected neurons. Similarly, although neurofilament aggregation is a pathological hallmark of several diseases^25,66^, previous *in vivo* studies have only reported perikaryal and axonal neurofilament aggregation.

Furthermore, we show that calpain activity is necessary for the nuclear accumulation of NFL and that either the intact tail domain of NFL or its proteolytic fragment localizes in the nuclei of primary neurons during oxidative injury *in vitro*. These results are consistent with previous studies that have shown calpain-mediated degradation of neurofilaments^41-43^. By overexpressing recombinant NFL domains, we further show that the tail domain of NFL accumulates in the nuclei of healthy neurons and ND7/23 cells. We confirmed this by staining with two anti-NFL antibodies and by observing a nuclear FLAG signal in injured neurons that expressed the recombinant calpain-insensitive NFL-FLAG. These results also show that the region between amino acid A461 and the C terminus is not necessary for the nuclear accumulation of the NFL tail domain to occur.

In addition, by utilizing DamID method ^56,57^, we demonstrate that the tail domain of NFL interacts with DNA in the nucleus, suggesting that NFL might influence gene expression. The idea that neurofilaments could regulate gene expression was proposed in the early days of the neurofilament research^67-69^. This hypothesis was supported by a series of studies that showed that neurofilaments and other intermediate filaments have nucleic acid-binding properties^70-73^ and an ability to interact with histones^74^. Another study on the mechanism of vimentin–DNA binding suggested that this interaction is mediated by intercalation between DNA base pairs^75^. Our results further support these general findings and demonstrate that NFL–DNA interaction also occurs in the nuclei of living cells, in addition to the test tube conditions utilized in the aforementioned studies. However, our results show that the tail domain of NFL, or its proteolytic fragment, localizes in the nucleus and interacts with the DNA upon injury. This is in contrast to the findings of Wang et al. who identified nucleic acid binding sites in the first half of the head domain of NFL^76^. For that reason, we examined if the N-terminal head domain localizes in the nuclei of nitric oxide-treated neurons, by utilizing an antibody that binds amino acids 6–25 of the NFL head domain. We observed no fluorescent signal in the nuclei of injured neurons, suggesting that the head domain of NFL does not localize there during injury. However, our NFL-domain overexpression experiments showed that the NFL head can be detected in the nucleus of neurons. Still, there was no clear nuclear accumulation of the NFL head domain such as that we observed for the NFL tail. In addition, as the NFL head domain contains only 93 amino acids (∼10 kDa), its presence in the nucleus might result from passive diffusion through nuclear pores. Nevertheless, in addition to the previously reported N-terminal DNA-binding site, our data indicate the existence of another DNA-binding site in the tail domain of NFL. These discrepancies suggest that the interaction between NFL and DNA is different in living cells, a situation far removed from the non-physiological test tube conditions utilized by Wang et al. However, in addition to the differences between NFL behavior in the test tube and *in vitro*, NFL response to the oxidative injury might also proceed differently *in vivo*. This is an important limitation of our study, that focuses solely on oxidative stress in neuronal cultures *in vitro*, which do not fully represent the complexity of an intact nervous system.

Finally, our results suggest that a proteolytic fragment rather than the full-length NFL might regulate the neuronal response to oxidative injury. Previous immunoblot studies have reported that proteolytic fragments are present in neurofilament preparations from brain, spinal cord, and peripheral nerves^41,77,78^. Additional studies have shown that neurofilament fragments have tubulin-binding properties, and that they can be taken up by both neuronal and glial cells where they influence microtubule dynamics^22,79,80^. Moreover, a recent study^81^ using mass spectrometry mapping identified a fragment containing amino acids 530-540 of the NFL tail domain in healthy brain tissue. Multiple NFL truncated species were identified in CSF, with varying amounts in Alzheimer’s disease, suggesting that these species could be differentially produced/secreted in physiological and pathophysiological conditions. Although this requires further investigation, altogether these studies suggest that degradation products of neurofilaments might have additional functions in the nervous system. Furthermore, it was shown that neurons lacking NFL are more susceptible to nitric oxide-induced injury and death^82^, whereas NFL knock-out mice have delayed axonal regeneration following spinal cord injury^83^. These findings support the possibility that NFL might play important roles in the neuronal response to injury. Our study suggests that these functions could be at least partially fulfilled by the accumulation of the NFL tail domain or its proteolytic fragment in the nucleus. Finally, as our results show that the tail domain of NFL interacts with the DNA, the accumulation of the NFL tail domain in the nucleus might be involved in the regulation of gene expression in response to oxidative injury.

## Materials and methods

### Cell culture

Primary mouse cortical neurons (Thermo Fisher Scientific, cat. no. A15586) and mouse hippocampal neurons (MHNs; Thermo Fisher Scientific, cat. no. A15587) from E17 C57BL/6 mice and primary rat cortical neurons (RCNs; Thermo Fisher Scientific, cat. no. A36512) from E18 Sprague Dawley rats were thawed and cultured according to the manufacturer’s recommendations in a B-27 Plus Neuronal Culture System (Thermo Fisher Scientific, cat. no. A3653401). The thawing and culturing medium (NB Plus +) consisted of Neurobasal Plus Medium containing 2% B-27 Plus Supplement and 1% penicillin–streptomycin (PS; Sigma−Aldrich, cat. no. P0781). After thawing, mouse cortical neurons were cultured for up to 20 days, and half the volume of medium was changed twice per week.

For experiments, neurons were seeded on eight-well Lab-Tek II chambered coverglasses (German #1.5 borosilicate glass; Thermo Fisher Scientific, cat. no. 155409) at a density of 70,000– 110,000 cells per well. Before seeding, the coverglasses were coated for 2 h at room temperature (RT) with a 20 μg/ml solution of poly-D-lysine (PDL; Sigma–Aldrich, cat. no. P6407) in sterile double distilled water (ddH_2_O). After coating, the coverglasses were washed three times with sterile ddH_2_O and allowed to dry inside a sterile hood at RT for at least 30 min. Then, warm NB Plus + medium was added to each well and the chambers were incubated for at least 30 min at 37 °C and 5% CO_2_ before neuron seeding.

ND7/23 cells (a mouse neuroblastoma × rat dorsal root ganglion neuron hybrid cell line; Sigma−Aldrich, ECACC 92090903), were cultured in high-glucose Dulbecco’s Modified Eagle’s Medium (DMEM; Thermo Fisher Scientific, cat. no. 41965062) containing 10% heat-inactivated fetal bovine serum (FBS; Thermo Fisher Scientific, cat. no. 10270106), 1% PS, 1% sodium pyruvate (Thermo Fisher Scientific, cat. no. 11360039) and 1% L-glutamine (Thermo Fisher Scientific, cat. no. 25030024). FBS was inactivated by incubation at 56 °C for 30 min. ND7/23 cells were passaged three times per week and used between passages 3 and 15.

ND7/23 cells were seeded on eight-well Lab-Tek II chambered coverglasses at a density of 25,000 cells per well. Prior to cell seeding, the coverglasses were coated with a 10 μg/ml solution of PDL in ddH_2_O for a minimum of 4 h at RT, washed three times with ddH_2_O, and allowed to dry inside a sterile hood.

IPSC-derived neurons were a kind gift from Dr. Stefan Hauser and Milena Korneck, DZNE Tuebingen, Germany. The cells were cultured and differentiated as previously described,^84,85^ and used for experiments on the 37th–39th day of differentiation.

### Compartmentalized microfluidic cultures

Silicone microfluidic devices with two compartments and a 450 μm microgroove barrier were purchased from Xona Microfluidics (cat. no. SND450) and prepared according to the manufacturer’s instructions with some modifications. Menzel coverslips (borosilicate glass #1, 0.13–0.16 mm thickness, 22×22 mm; Fisher Scientific, cat. no. 17214904) were cleaned by sonication for 30 min and subsequent washing in 70% ethanol for 1 h. Then, coverslips were thoroughly washed with sterile ddH_2_O, incubated in sterile ddH_2_O for 1 h, and allowed to dry inside a sterile hood at RT. Coverslips were then coated with 100 μg/ml PDL solution (Thermo Fisher Scientific, cat. no. A3890401) for 4 h at 37 °C and 5% CO_2_, washed three times with sterile ddH_2_O, and kept submerged in sterile ddH_2_O overnight (ON) at 4 °C. On the following day, coverslips were washed with sterile ddH_2_O and allowed to dry inside a sterile hood at RT. The microfluidic devices were immersed for 1 h in 70% ethanol, washed with sterile ddH_2_O, then incubated for 1 h in sterile ddH_2_O, and kept inside a sterile hood at RT until completely dry. The microfluidic devices were assembled on top of the coated coverslips, loaded with warm NB Plus + medium and incubated ON at 37 °C and 5% CO_2_. On the following day, the channels of the device were checked under a microscope for the presence of air bubbles. If present, bubbles were removed by gently pipetting medium through the channel. Mouse cortical neurons were loaded into one compartment of each device at a density of 100,000–150,000 neurons per device, according to the manufacturer’s instructions. During culturing, half the volume of medium was exchanged every two days. Fluidic pressure was maintained by adding twice the volume of medium into the somatic compartment than in the axonal compartment.

### Chemicals, antibodies and fluorescent dyes

To study the effects of oxidative stress in neurons we used the following chemicals: sulfo NONOate (negative control of nitric oxide-induced stress; Cayman Chemical, cat. no. 83300), spermine NONOate (Cayman Chemical, cat. no. 82150), hydrogen peroxide (Sigma−Aldrich, cat. no. H1009), CCCP (Sigma−Aldrich, cat. no. C2759), and monosodium glutamate (Sigma−Aldrich, cat. no. G5889).

For protease inhibition we used calpain inhibitor III (Cayman Chemical, cat. no. 14283), EST (Sigma−Aldrich, cat. no. 330005), and emricasan (Cayman Chemical, cat. no. 22204).

For immunocytochemical staining we used the following primary and secondary antibodies: mouse anti-NFL antibody clone DA2 (Merck Millipore, cat. no. MAB1615), mouse anti-NFL antibody clone F-12 (Santa Cruz Biotechnology, cat. no. sc-390732), rabbit anti-NFL antibody clone EPR22035-112 (Abcam, cat. no. ab223343), rabbit anti-NFL antibody clone C28E10 (Cell Signaling Technology, cat. no. 2837), rabbit anti-NFM antibody (BioLegend, cat. no. 841001), mouse anti-nonphosphorylated NFH antibody (SMI-32P; BioLegend, cat. no. 801701), rabbit anti-NeuN antibody (Merck Millipore, cat. no. ABN78), rabbit anti-FLAG antibody (Merck Millipore, cat. no. F7425), goat anti-mouse Alexa Fluor(AF) 488 Plus (Thermo Fisher Scientific, cat. no. A32723), goat anti-mouse AF647 Plus (Thermo Fisher Scientific, cat. no. A32728), goat anti-rabbit AF555 (Thermo Fisher Scientific, cat. no. A21429), and goat anti-rabbit AF647 Plus (Thermo Fisher Scientific, cat. no. A32733). For the staining of nuclei, we used the green fluorescent nucleic acid stain SYTO16 (Thermo Fisher Scientific, cat. no. S7578).

Anti-NFL antibody clone DA2 (Merck Millipore cat. no. MAB1615) was used for immunostaining in all experiments except for that yielding the data shown in Supplementary Fig. 2. The anti-NFL antibodies clone F-12 (Santa Cruz Biotechnology, cat. no. sc-390732), EPR22035-112 (Abcam, cat. no. ab223343), and C28E10 (Cell Signaling Technology, cat. no. 2837) were used for immunostaining of NFL only for the experiment yielding the data shown in Supplementary Fig. 2.

### Treatments

Treatments of mouse cortical neurons, MHNs, and RCNs with the control compound sulfo NONOate and the nitric oxide-producing spermine NONOate were performed on either day *in vitro* (DIV) 10 or DIV 12. First, the culture medium was aspirated from the wells and replaced by fresh warm NB Plus +, followed by pre-incubation for 4 h at 37 °C and 5% CO_2_. Then, diluted solutions of NONOates were prepared in fresh warm NB Plus +, at concentrations 2× greater than that desired. These solutions were then added to wells such that the volumes of the treatment and the medium already in the well were in a 1:1 ratio, thereby ensuring the final desired concentration of NONOate. Neurons were incubated with the NONOates for the desired amount of time (usually 3 h) at 37 °C and 5% CO_2_, and then either fixed for immunostaining or stained with annexin V and propidium iodide. For experiments involving protease inhibition, calpain and caspase inhibitors were added to the medium 1 h before injury. The working concentrations of calpain inhibitor III and EST were 100 μM and that of emricasan was 20 μM.

Mouse cortical neurons grown in microfluidic devices were treated with sulfo NONOate and spermine NONOate in a similar way. Four hours before injury, the medium was aspirated from the compartments undergoing treatment, and replaced with fresh warm NB Plus +. After 4 h of pre-incubation, the medium was aspirated from these compartments and mixed at a 1:1 ratio with fresh warm NB Plus +. NONOates were added to these mixtures at a final concentration of 1 mM, and these treatments were added back to the wells. After incubation for 3 h at 37 °C and 5% CO_2_, the medium was removed from wells, the device was removed from the coverslip, and neurons were fixed for immunocytochemical staining.

IPSC-derived human neurons were treated on the 37–39^th^ day of differentiation with 2 mM sulfo NONOate or spermine NONOate. First, the 3N culturing medium (1:1 DMEMF12/N2:Neurobasal/B-27)^84^ was aspirated and replaced with 3N medium without additives (“stressing” medium), and cells were incubated for 4 h at 37 °C and 5% CO_2_. After this pre-incubation, treatments were performed by aspirating the medium from the wells and replacing it with a 2 mM solution of the NONOate in the stressing medium. After 3 h, medium was aspirated once more and replaced with fresh 2 mM NONOate solution. After incubation for a further 2 h, neurons were fixed for immunocytochemical staining.

Treatment of mouse cortical neurons with 25 μM CCCP was done on DIV 14–16 (3 h), treatment with 100 μM monosodium glutamate was performed on DIV 14–20 (6 h), and treatment with 1 mM H_2_O_2_ was done at DIV 10 (3 h). These treatments were performed according to the same protocol described for NONOate treatments.

### Constructs and cloning

The cloning of plasmids encoding NFL-mGFP and NFL-FLAG was described previously^86^. Head (amino acids 1–93), rod (amino acids 94–397), and tail (amino acids 398–543) domains of NFL were amplified by PCR from the pCMV-NFL-mGFP plasmid. The sequence encoding full-length NFL was excised from the pCMV-NFL-FLAG plasmid with the enzymes HindIII and BamHI, and NFL domains containing the same restriction sites were ligated with this backbone after the CMV promoter.

Calpain-insensitive constructs were cloned by PCR amplification of NFL amino acids 1–461 from the vector pCMV-NFL-FLAG with the primers that contained either HindIII and BamHI sites (for cloning with the GFP) or HindIII and NotI sites (for cloning with the FLAG tag). The NFL1–461 sequence with HindIII and BamHI sites was cloned into the mEGFP-N1 plasmid previously cut with the corresponding enzymes, to yield the NFL(ΔA461–D543)-GFP construct. To obtain the NFL(ΔA461–D543)-FLAG construct, the sequence encoding the full-length NFL was excised from the pCMV-NFL-FLAG plasmid with the enzymes HindIII and NotI, and the NFL1–461 sequence with the same sites was ligated into the resulting backbone.

The pmNFM plasmid containing cDNA encoding for NFM was a gift from Anthony Brown (Addgene plasmid #83126; http://n2t.net/addgene:83126; RRID: Addgene_83126)^87^.

Plasmids encoding *Escherichia coli* DNA adenine methyltransferase (Dam) and GFP-^m6^A-Tracer were a kind gift from Dr. Bas van Steensel (Netherlands Cancer Institute, Amsterdam, Netherlands). To generate a control plasmid, the gene encoding the Dam was amplified by PCR with primers containing HindIII and EcoRI restriction sites, and subsequently cloned into an empty pcDNA3.1/Zeo(+) plasmid backbone. Genes encoding either full-length NFL or the NFL tail domain were amplified by PCR with primers containing NheI and HindIII restriction sites. They were then cloned in-frame into the pcDNA3.1/Zeo(+)-Dam plasmid to generate pcDNA3.1/Zeo(+)-NFL-Dam and pcDNA3.1/Zeo(+)-NFL(tail)-Dam constructs. The plasmid encoding GFP-^m6^A-Tracer was used without modification.

### Transfections

Mouse cortical neurons were transfected as described previously at DIV 8^86,88^. In brief, mouse cortical neurons were transfected using the Lipofectamine 2000 transfection reagent (Thermo Fisher Scientific, cat. no. 11668027), with a DNA/Lipofectamine 2000 ratio of 1 μg:2.4 μl. Plasmid DNA and Lipofectamine 2000 were added to Neurobasal Plus Medium containing 1% PS. For experiments with the full-length NFL and NFL domains, mouse cortical neurons were transfected with either 0.25 μg of NFL plasmid alone, or with 0.25 μg of NFL plasmid and 0.125 μg of NFM per well. For experiments with calpain-insensitive NFL constructs, mouse cortical neurons were transfected with 0.5 μg of NFL construct and 0.25 μg of NFM per well. For DamID experiments, mouse cortical neurons were transfected with 0.25 μg of NFL-Dam or NFL(tail)-Dam plasmid with 0.125 μg of NFM plasmid and 0.25 μg of GFP-^m6^A-Tracer per well. Eight-well Lab-Tek chambered slides were used. After preparation, the transfection solutions were mixed at a 1:1 ratio with Neurobasal Plus Medium containing 1% PS and 4% B-27 Plus, to obtain the 2% B-27 Plus content recommended for the culturing of mouse cortical neurons. Transfection mixtures were then incubated for 5 min at 37 °C and 5% CO_2_. The culturing medium was aspirated from the neurons and retained for use as a conditioned medium. Immediately after, transfection solutions were added dropwise to the neurons, which were then incubated for 6 h at 37 °C and 5% CO_2_. Transfection solutions were then aspirated from the wells and replaced with a 1:1 mixture of fresh warm NB Plus + and conditioned medium. Mouse cortical neurons were incubated for 3 days at 37 °C and 5% CO_2_, and then either fixed or treated with NONOates and then fixed, for immunocytochemical staining. Mouse cortical neurons expressing NFL-GFP constructs were imaged live three days post-transfection during the NONOate treatment.

ND7/23 cells were transfected 14–20 h after seeding using Lipofectamine 2000 according to a slightly modified manufacturer’s protocol. The same DNA/Lipofectamine ratio and amounts of DNA in each well of an eight-well Lab-Tek chambered slide were used as for the transfection of mouse cortical neurons with full-length NFL and NFL domains. For DamID experiments, ND7/23 cells were transfected with 0.25 μg of full-length NFL-Dam or NFL(tail)-Dam plasmid and 0.25 μg of GFP-^m6^A-Tracer with or without 0.125 μg of NFM plasmid per well. Control cells were transfected with 0.25 μg of Dam with 0.25 μg of GFP-^m6^A-Tracer, or with 0.25 μg GFP-^m6^A-Tracer alone, per well. DNA and Lipofectamine 2000 were added to Opti-MEM I Reduced Serum Medium (Thermo Fisher Scientific, cat. no. 31985062). Prepared transfection mixtures were added on top of the ND7/23 culturing medium, and incubated for 6 h at 37 °C and 5% CO_2_. The culturing medium was then exchanged, cells were incubated ON at 37 °C and 5% CO_2_, and fixed for immunocytochemical staining on the following day.

### Immunocytochemical staining

In all experiments, neurons were fixed for 15 min at RT with 4% electron microscopy grade paraformaldehyde (PFA; Electron Microscopy Sciences, cat. no. 15710) diluted in PIPES–EGTA– magnesium (PEM) buffer (80 mM PIPES, 2 mM MgCl_2_, 5 mM EGTA, pH 6.8). After fixation, neurons were rinsed three times with 0.01 M phosphate-buffered saline (PBS; 137 mM NaCl, 10 mM Na_2_HPO_4_, 1.8 mM KH_2_PO_4_, 2.7 mM KCl, pH 7.4) and blocked for 1 h at RT with a serum containing 3% bovine serum albumin (BSA; Sigma−Aldrich, cat. no. A9647), 10% goat serum (Thermo Fisher Scientific, cat. no. 16210072), and a 0.2% solution of Triton X-100 (Sigma−Aldrich, cat. no. X100) in PBS. PBS was replaced with Tris-buffered saline (TBS; 20 mM Tris, 150 mM NaCl, pH 7.6) for the staining of nonphosphorylated NFH. Primary antibodies diluted in the blocking serum were then added to neurons and incubated for either 1 h at RT or ON at 4 °C. Neurons were then washed three times with PBS (5 min per wash), and incubated for 1 h at RT with a blocking serum containing secondary antibodies and the nuclear dye SYTO16. After this incubation step, the neurons were washed three times with PBS (5 min per wash) and either imaged immediately or kept at 4 °C until imaging.

ND7/23 cells were immunostained according to the same protocol described for neurons, with some modifications. The medium was aspirated from the wells, and cells were rinsed once with PBS and fixed for 15 min at RT with 4% PFA (Sigma−Aldrich, cat. no. 158127) in 0.1 M phosphate buffer. After fixation, cells were washed three times with PBS (5 min per wash) and immunostained by following the protocol described above.

Primary antibodies were used at the following dilution ratios: mouse anti-NFL antibody clone DA2, 1:300 or 1:500; mouse anti-NFL antibody clone F-12, 1:200; rabbit anti-NFL antibody clone EPR22035-112, 1:500; rabbit anti-NFL antibody clone C28E10, 1:500; rabbit anti-NFM antibody, 1:500; mouse anti-non-phosphorylated NFH antibody, 1:300; rabbit anti-FLAG antibody, 1:1000; rabbit anti-NeuN antibody, 1:500. All secondary antibodies were used at a 1:500 dilution, and SYTO16 dye was used at 1:1000.

### Cell death assays

For the terminal deoxynucleotidyl transferase dUTP nick end labeling (TUNEL) assay, mouse cortical neurons were treated at DIV 9–12 with 250 μM sulfo NONOate or 250 μM spermine NONOate for 3 h at 37 °C and 5% CO_2_, and then fixed with 4% electron microscopy grade PFA diluted in PEM buffer, as described above. Mouse cortical neurons were then permeabilized with 0.25% Triton X-100 in PBS for 20 min at RT, and positive-control wells were treated with 9.6 units of DNase I (Sigma−Aldrich, cat. no. D5025) per well. DNase I digestion was performed for 30 min on a heating block set to 37 °C. The neurons were then rinsed three times with molecular biology water (Sigma−Aldrich, cat. no. W4502) and TUNEL staining was performed by following the protocol provided with the kit (Thermo Fisher Scientific, cat. no. C10617). After the TUNEL assay was completed, cells were blocked and immunostained with anti-NFL antibody clone DA2, as described above.

For the annexin V/propidium iodide assay (using a staining kit from Sigma−Aldrich, cat. no. 11858777001), mouse cortical neurons were treated at DIV 12 with either 500 μM sulfo NONOate or 500 μM spermine NONOate for 2.5 h at 37 °C and 5% CO_2_. For the positive control of apoptosis, mouse cortical neurons were treated for 2.5 h with 500 μM sulfo NONOate and 10 μM staurosporine (Sigma−Aldrich, cat. no. S4400). For the positive control of necrosis, mouse cortical neurons were treated with 500 μM sulfo NONOate for 2.5 h and 0.1% saponin (Roth, cat. no. 9622) for 15 min. After treatment, mouse cortical neurons were labeled with annexin V and propidium iodide according to the protocol provided by the manufacturer. Mouse cortical neurons were then imaged live at 37 °C with a widefield epifluorescent microscope (described below). Live imaging was performed in Hibernate E medium (Brain Bits LLC, cat. no. HELF) supplemented with 2% B-27 Plus and 1% PS.

### Widefield imaging of fixed and live cells

Live- and fixed-cell widefield imaging was performed on an inverted Nikon Eclipse Ti2-E microscope (Nikon Instruments), controlled by NIS-Elements AR software (Nikon Instruments). The microscope features a XY-motorized stage, Perfect Focus System, and Apo 60× (NA 1.4, oil) and HP Apo TIRF 100× (NA 1.49, oil) objectives. The excitation light was provided by a fluorescent lamp (Lumencor Sola SE II), and filtered through 488 (AHF; EX 482/18; DM R488; BA 525/45), 561 (AHF; EX 561/14; DM R561; BA 609/54), and Cy5 (AHF; EX 628/40; DM660; BA 692/40) filter cubes. Emitted light was imaged with ORCA-Flash 4.0 sCMOS camera (Hamamatsu Photonics). Images were acquired at a 16-bit depth, 1024×1024 pixels, and pixel size of either 0.27 or 0.16 μm.

For the live-cell imaging of annexin V and PI-labeled cells, a microscope cage incubator with dark panels (Okolab, Naples, Italy) was pre-warmed and maintained at 37 °C during the imaging. An H201-T-UNIT-BL temperature unit (Okolab) was used to control the temperature. Live-cell imaging was performed in Hibernate E medium containing 2% B-27 Plus and 1% PS.

Live-cell imaging of neurons expressing NFL-GFP and NFL(ΔA461–D543)-GFP was performed by using the incubator and the heating unit described above. The incubator was pre-warmed to 37 °C and neurons in artificial cerebro-spinal fluid (aCSF) containing 2% B-27 Plus and 1% PS were placed on the stage. In each well, 10 neurons expressing either NFL-GFP or NFL(ΔA461–D543)-GFP were selected, and the *xyz* coordinates of each field of view were saved in NIS-Elements AR software. After the acquisition of the first set of images, NONOate solutions were prepared in aCSF (containing 2% B-27 Plus and 1% PS), to a double concentration of 1 mM. Diluted NONOates were added to wells at a 1:1 ratio with the volume of medium already present to achieve a final concentration of 0.5 mM. Images were acquired immediately after the addition of the NONOate, and then every 10 min until approximately 1 h after treatment began.

### Image processing

The brightness and contrast of raw widefield images were linearly adjusted in Fiji software^89^. Images were converted to 8-bit depth and saved as TIFF files. The analyzed images were imported into Adobe Illustrator for arrangement into the figures in this article.

Raw 16-bit images were used for the measurement of fluorescence intensity line profiles in Fiji and the *xy* coordinates of line profiles were exported to Microsoft Excel. In Excel, the *xy* coordinates were used to generate graphs that were then imported into Adobe Illustrator for the preparation of figures.

### Nuclear NFL and GFP fluorescence intensity measurements

To measure the fluorescence intensity of NFL in the nucleus after treatment with calpain or caspase inhibitor and nitric oxide (Fig. 4b), images were acquired on the widefield microscope described above. Three independent experiments were performed, with 10 images acquired per experiment per condition. Images were analyzed in Fiji in a semi-automated manner and done blindly to avoid bias. Images of NFL and the corresponding images of SYTO16 and bright-field images were opened in Fiji. Regions of interest (ROIs) for the measurement were defined by applying the automatic Otsu dark thresholding algorithm to the SYTO16 images. These ROIs were subsequently applied on bright-field images and all ROIs that were not nuclei were discarded. In addition, all ROIs containing high-intensity out-of-focus fluorescence originating from the surrounding axons or dendrites were excluded from the analysis. In the next step, selected ROIs were applied on the NFL images and used for measurements of area, mean intensity, integrated density, and raw integrated density. In addition to nuclear ROIs, a rectangular ROI was used to measure background values for each image. These background ROIs were placed in a region free of cells. Corrected total cell fluorescence (CTCF) was calculated using the measured values and the formula: CTCF = raw integrated density − (area × mean fluorescence of the image background). The calculated CTCF values were divided by the areas of the corresponding nuclei and the resulting values were used to compare NFL florescence intensities between the groups.

The analysis of nuclear GFP fluorescence intensity (Fig. 8f) was performed similarly. Images were acquired on the widefield microscope. Two independent experiments were performed, with 10 images acquired per experiment per condition. Images were analyzed in Fiji in a semi-automated manner and done blindly to avoid bias. GFP images were opened in Fiji and ROIs were defined by applying the automatic Li dark thresholding algorithm. As described above, all ROIs that were not nuclei were removed and a rectangular ROI was placed in a region free of cells to measure background values for each image. In all these ROIs we measured area, mean intensity, integrated density, and raw integrated density, and calculated CTCF values. CTCF values were divided by the area of each ROI and these values were later used for the statistical analysis.

### NeuN- and NFL-positive cell counting

For the counting of NeuN- and NFL-positive cells (Supplementary Fig. 1b), images of neurons stained with anti-NeuN and anti-NFL antibodies were acquired on the widefield microscope described above. Cells with NeuN+ staining were identified in Fiji by applying the automatic Default dark thresholding algorithm. Subsequently, cells were counted by transforming the image into a binary mask, applying watershed segmentation to separate adjacent cells, and automatic counting of all cells with a radius greater than 30. Cells with NFL+ nuclei were counted manually, and the percentage of NFL+ cells from all NeuN+ neurons was calculated in Excel.

### Statistical analysis

Statistical analyses of the data presented in Figs. 4 and 8 were performed in IBM SPSS Statistics Version 28 (Armonk, New York, USA). First, the normality of data distribution was evaluated with the Kolmogorov–Smirnov test, which showed that the data were not normally distributed. Accordingly, the non-parametric Kruskal–Wallis test was applied to the data and indicated that there is an overall significant difference between the groups. To determine which groups were significantly different, a subsequent pairwise comparison was carried out with Dunn–Bonferroni correction for multiple comparisons. Results of these comparisons are shown in Supplementary tables 1 and 4.

## Acknowledgements

We thank Katja Widmaier and Het Mehta for their excellent technical assistance and all the members of the Nikić-Spiegel group for their support. We are grateful to Dr. Stefan Hauser and Milena Korneck for the gift of differentiated IPSC-derived human neurons, to Dr. Bas van Steensel for the gift of plasmids containing Dam methylase and GFP-^m6^A-Tracer, and to Dr. Anthony Brown for the gift of the plasmid containing NFM, which was obtained through Addgene. We also thank George Philippos and Nevena Stajković for their help with the cloning of plasmids containing NFL domains and Dam methylase. This study was supported by the Emmy Noether Programme (project number 317530061, to I.N.-S.) of the German Research Foundation (DFG) and the Werner Reichardt Centre for Integrative Neuroscience (Ministry of Science Baden-Württemberg).

## Author contributions

A.A. performed the experiments and collected the data. A.A. and I.N.-S. designed the experiments, analyzed the data, prepared the figures, and wrote the manuscript. I.N.-S. conceived and supervised the project.

