## Supplementary Information for "The tail domain of neurofilament light chain accumulates in neuronal nuclei during oxidative injury"

Supplementary Figures 1-10

Supplementary Tables 1-4

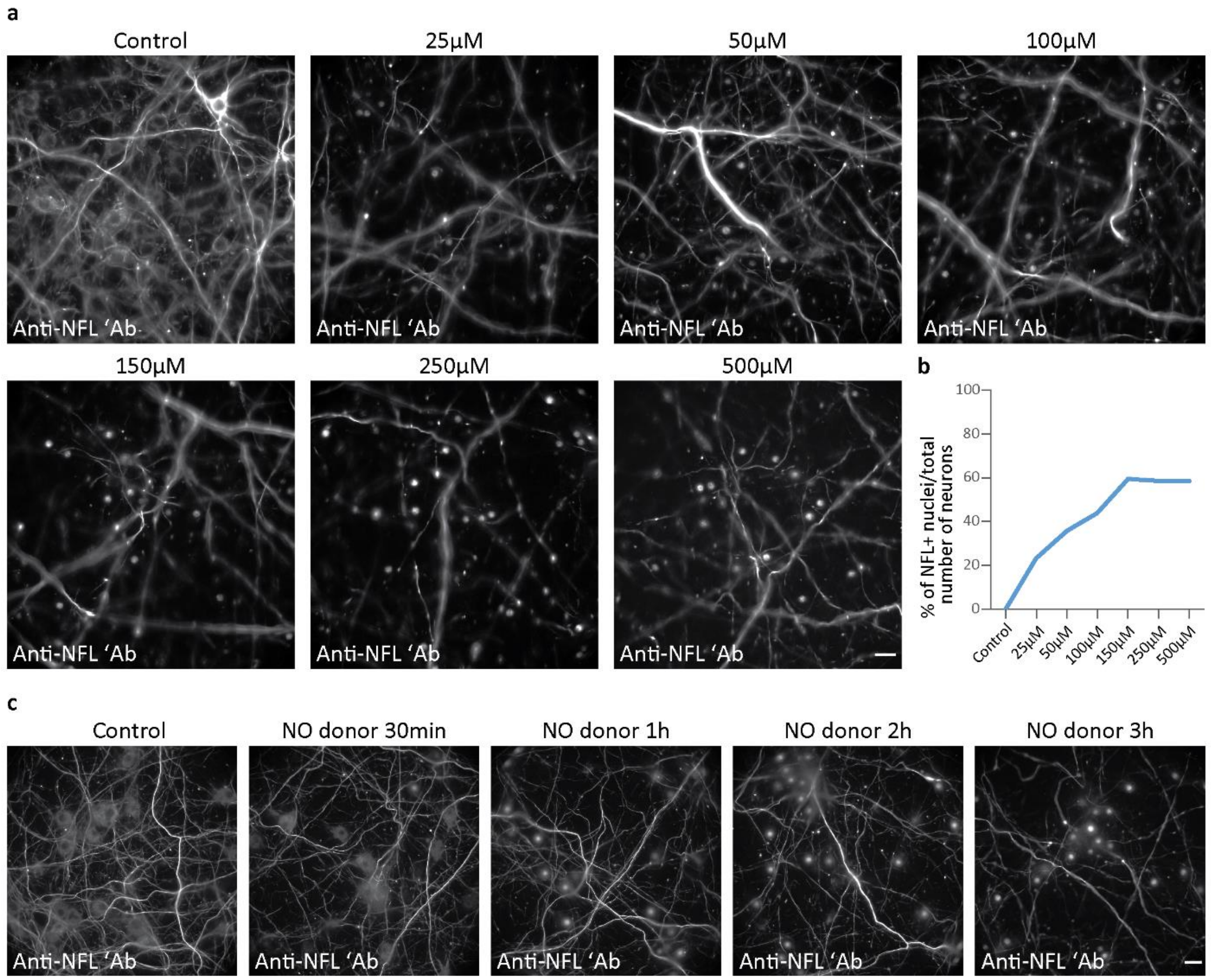


**Supplementary Fig. 1 | Localization of neurofilament light chain (NFL) in mouse cortical neurons (MCNs) after treatment with various concentrations of nitric oxide and incubation for different periods of time.** **a:** MCNs were treated at day *in vitro* (DIV) 10 for 3h with either 500 µM sulfo NONOate (control) or with varying concentrations of the nitric oxide donor spermine NONOate. After treatment, neurons were fixed, stained with anti-NFL and anti-NeuN antibodies, and imaged with widefield microscopy. **b:** Quantification of NFL-positive (NFL+) nuclei in MCNs treated and stained as described in **a**. The number of NFL+ nuclei is expressed as the percentage of the total number of neurons identified by NeuN staining. **c:** MCNs were treated at DIV 12 with 250 µM sulfo NONOate for 3 h (control) or 250 µM spermine NONOate (NO donor) for different periods of time. After the treatment, neurons were fixed, stained with anti-NFL antibody, and imaged with widefield microscopy. Scale bars: 20 µm.

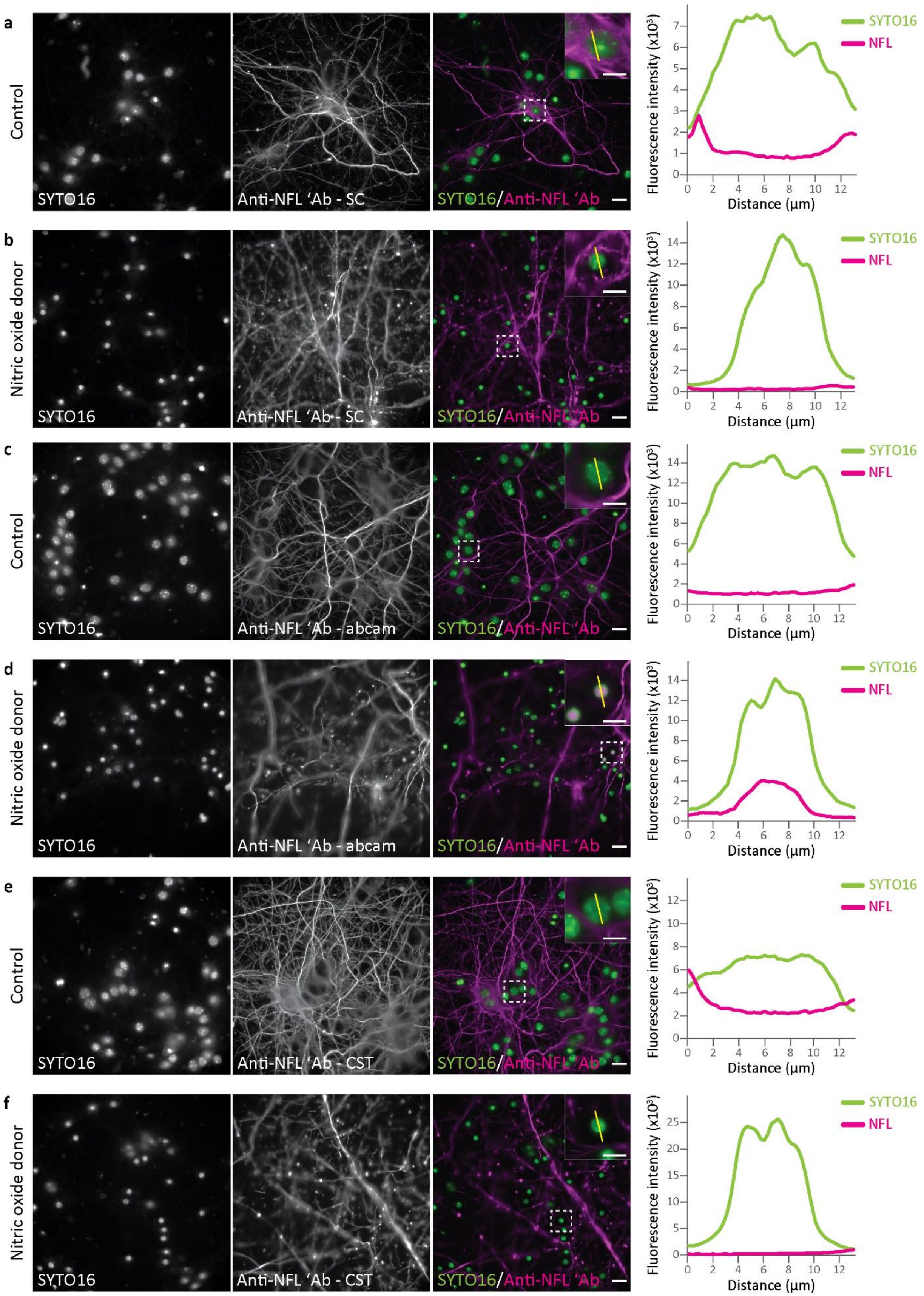
a
Control
SYTO16
Anti-NFL 'Ab - SC
SYTO16/Anti-NFL 'Ab
Fluorescence intensity (x10³)
Distance (µm)
SYTO16
NFL
b
Nitric oxide donor
SYTO16
Anti-NFL 'Ab - SC
SYTO16/Anti-NFL 'Ab
c
Control
SYTO16
Anti-NFL 'Ab - abcam
SYTO16/Anti-NFL 'Ab
d
Nitric oxide donor
SYTO16
Anti-NFL 'Ab - abcam
SYTO16/Anti-NFL 'Ab
e
Control
SYTO16
Anti-NFL 'Ab - CST
SYTO16/Anti-NFL 'Ab
f
Nitric oxide donor
SYTO16
Anti-NFL 'Ab - CST
SYTO16/Anti-NFL 'Ab

**Supplementary Fig. 2 | Immunocytochemical staining of NFL with three additional anti-NFL antibodies in control and injured neurons. a–f:** MCNs were treated at DIV 12 for 3h with either 500 µM sulfo NONOate (control; **a,c,e**) or 500 µM donor spermine NONOate (nitric oxide donor; **b,d,f**). After treatment, neurons were fixed and stained with anti-NFL antibody clones F-12 from Santa Cruz Biotechnology (SC; **a,b**), EPR22035-112 from Abcam (**c,d**), or C28E10 from Cell Signaling Technology (CST; **e,f**), and the nuclear dye SYTO16. After immunostaining, neurons were imaged with widefield microscopy. Graphs on the right show line profile fluorescence intensity measurements for both SYTO16 and anti-NFL signals. Fluorescence intensities are plotted as absolute gray values of raw 16-bit depth images. The regions of interest outlined by dashed boxes contain the nuclei used for the line profile measurement, expanded views of which are shown inset. Scale bars: 20 µm (**a–f**), 10 µm (images inset in **a–f**).

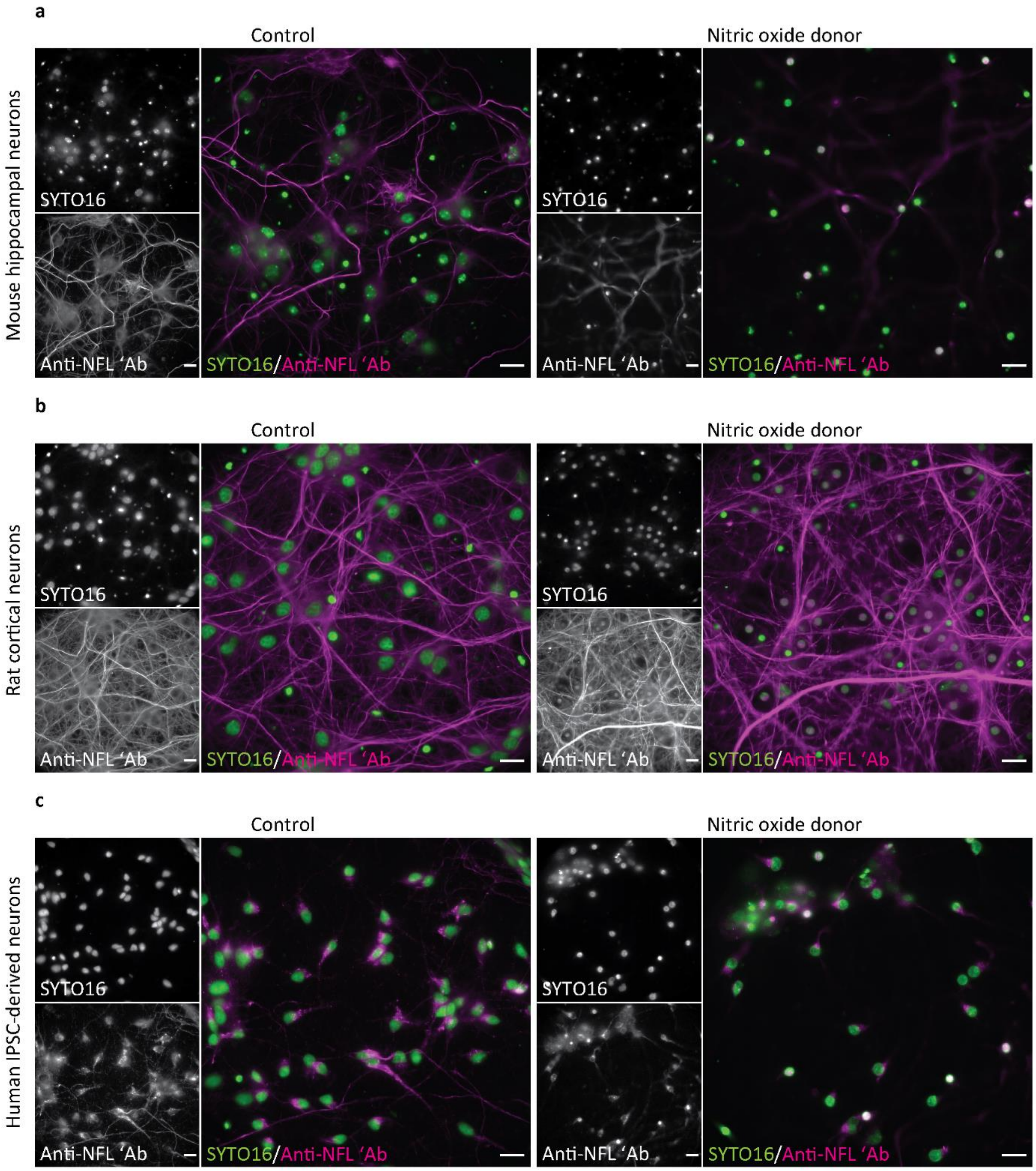


**Supplementary Fig. 3 | Localization of NFL after nitric-oxide-induced injury of different types of neurons. a:** Mouse hippocampal neurons were treated at DIV 12 for 3h with either 250 µM sulfo NONOate (control) or 250 µM spermine NONOate (nitric oxide donor). **b:** Rat cortical neurons were treated at DIV 12 for 3h with either 500 µM sulfo NONOate (control) or 500 µM spermine NONOate (nitric oxide donor). **c:** Human induced pluripotent stem cell (IPSC)-derived neurons were treated for 5h with either 2 mM sulfo NONOate (control) or 2 mM spermine NONOate (nitric oxide donor). After treatment, neurons were fixed, stained with anti-NFL antibody and nuclear dye SYTO16, and imaged with widefield microscopy. Scale bars: 20 µm.

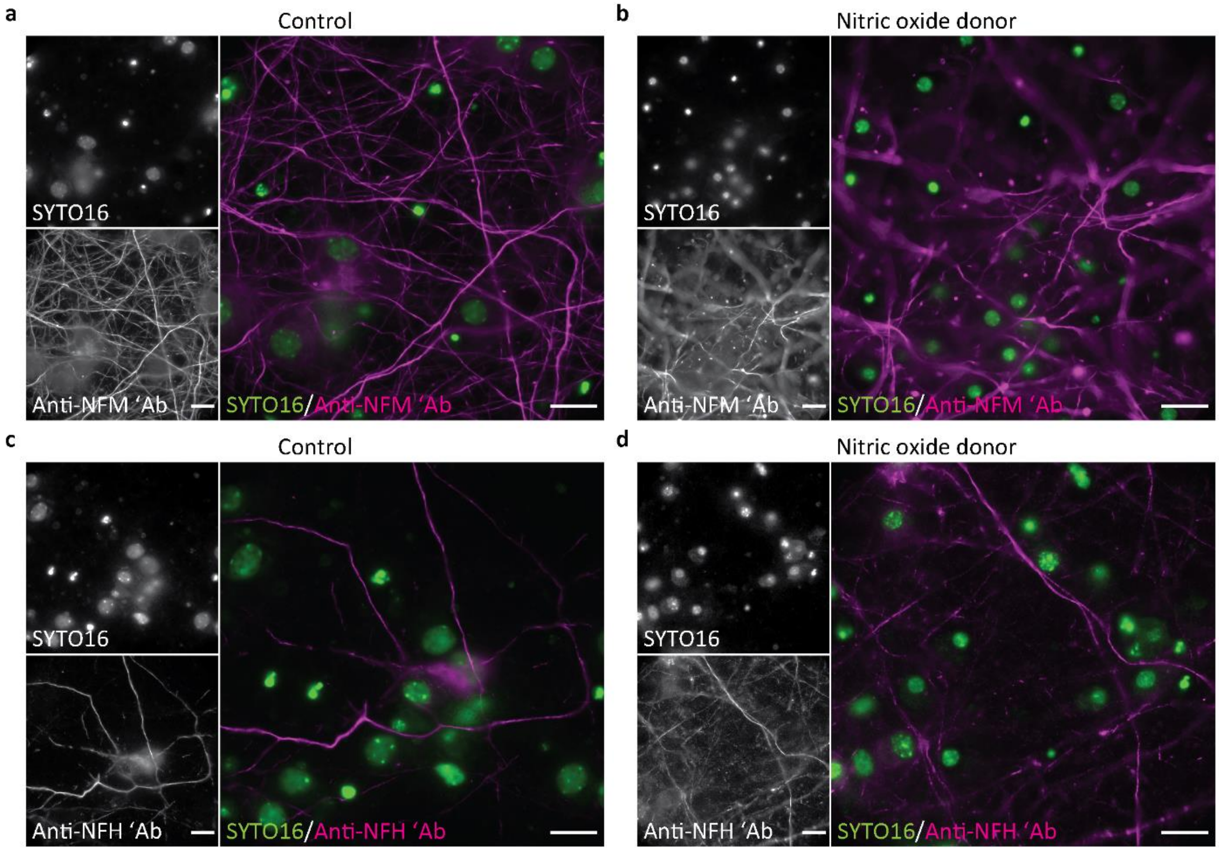


**Supplementary Fig. 4 | Labeling of other neurofilament subunits in control and injured neurons. a–d:** MCNs were treated for 3 h with either 500 µM sulfo NONOate (control; **a,c**) or 500 µM spermine NONOate (nitric oxide donor; **b,d**). After treatment, neurons were fixed and labeled with the nuclear dye SYTO16 and either anti-neurofilament medium chain (NFM; **a,b**) or anti-nonphosphorylated neurofilament heavy chain (NFH; **c,d**). Scale bars: 20 µm.

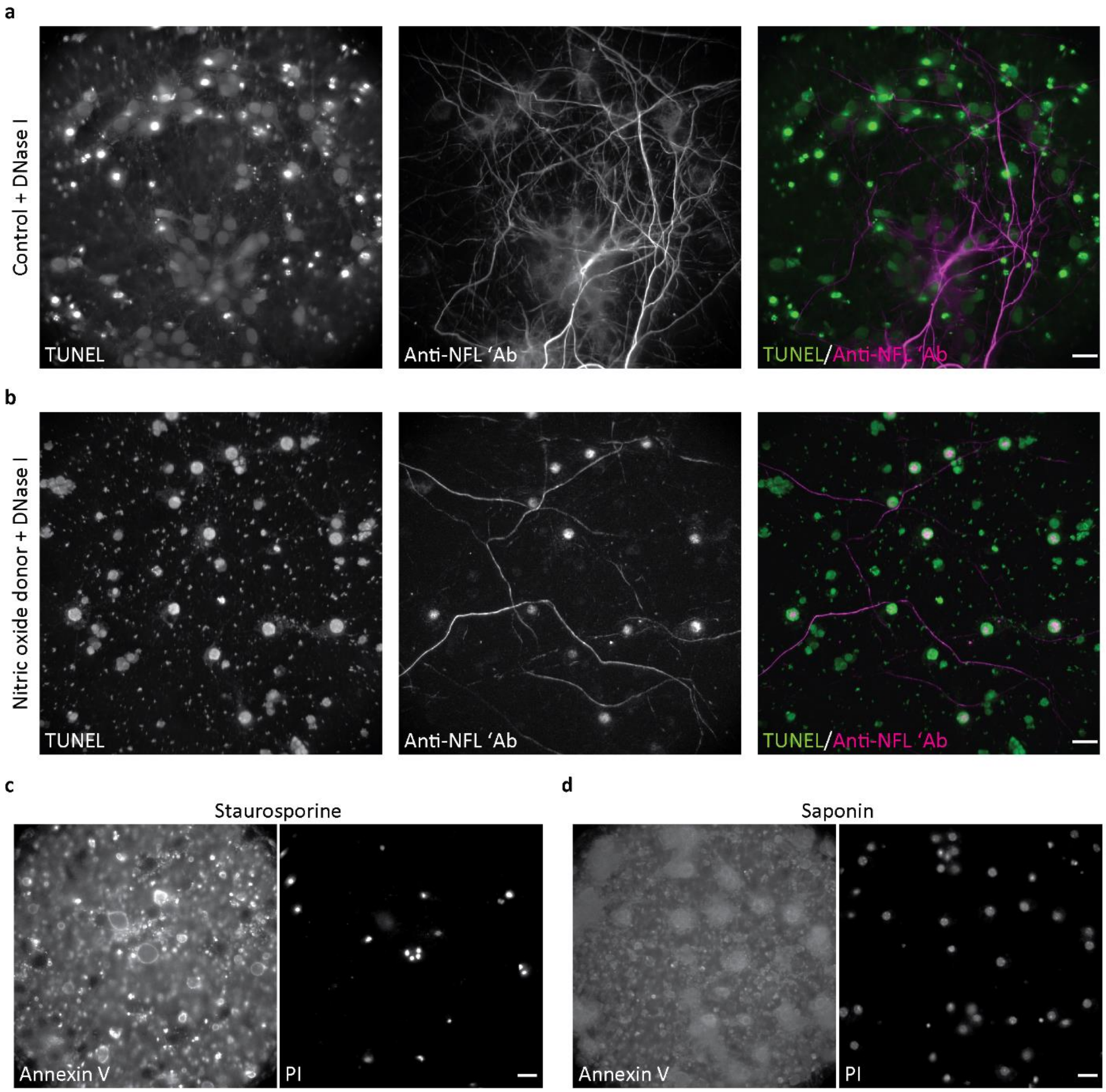


**Supplementary Fig. 5 | Positive control staining for cell death assays. a,b:** MCNs were treated at DIV 10 for 3 h with either 250 µM sulfo NONOate (control; **a**) or 250 µM spermine NONOate (nitric oxide donor; **b**), then fixed, permeabilized, and incubated with DNase I. Neurons were then labeled in a terminal deoxynucleotidyl transferase dUTP nick end labeling (TUNEL) assay to detect late-stage apoptosis, and with anti-NFL antibody. Images were acquired on a widefield microscope. **c,d**: MCNs were treated at DIV 12 with 500 µM sulfo NONOate for 2.5 h and with either 10 µM staurosporine for 2.5 h (**c**) or 0.1% saponin for 15 min (**d**). Neurons were then labeled with annexin V and propidium iodide (PI) for the detection of early-stage apoptosis or necrosis, respectively, and imaged live with widefield microscopy. Scale bars: 20 µm.

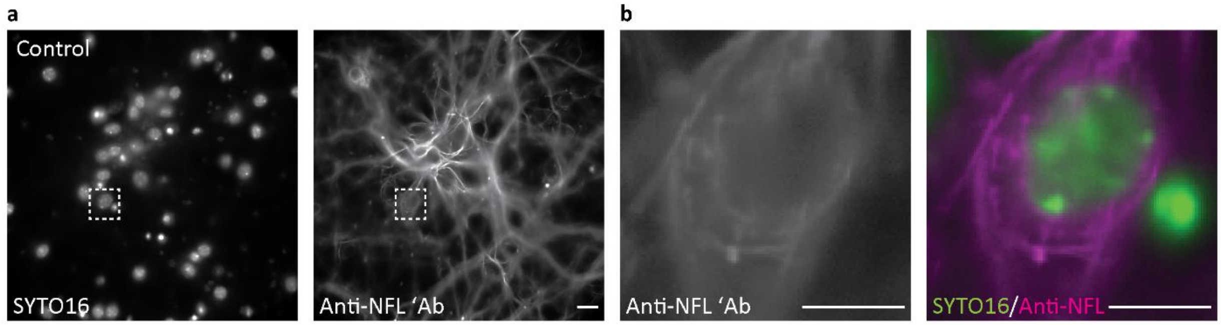


**Supplementary Fig. 6 | Representative image of control neurons used for nuclear NFL fluorescence intensity quantification. a,b:** DIV 12 MCNs were treated for 3 h with 500 µM sulfo NONOate. Neurons were then fixed, stained with anti-NFL antibody and the nuclear dye SYTO16, and imaged with widefield microscopy. **a:** Representative image of sulfo NONOate-treated neurons (control), which were used to quantify the nuclear NFL fluorescence intensity shown in Fig. 4. **b**: Images of a neuron from the dashed-box region in **a** with an out-of-focus anti-NFL signal in the nucleus. Scale bars: 20 µm (**a**), 10 µm (**b**).

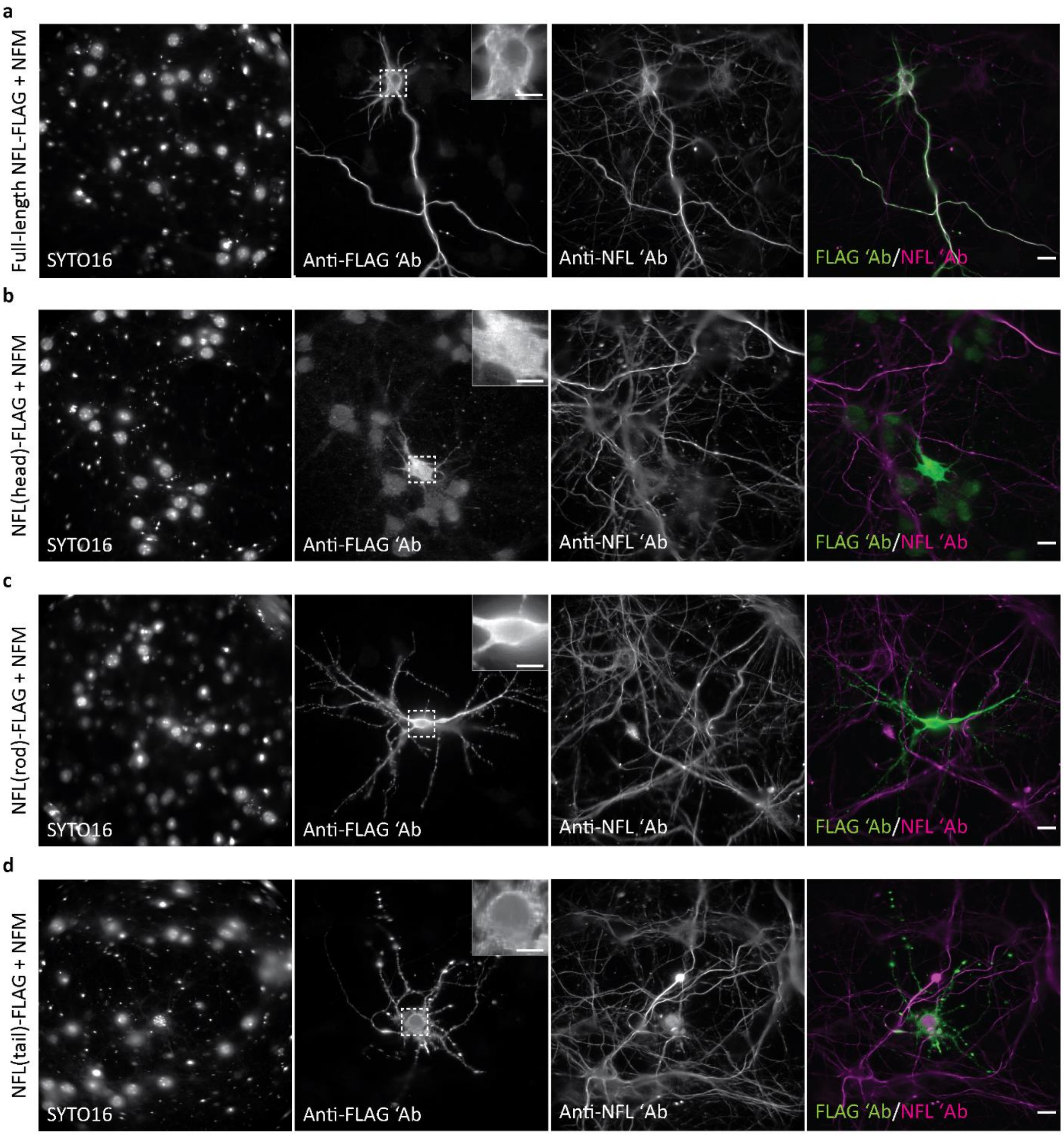


**Supplementary Fig. 7 | Localization of recombinant full-length NFL and NFL domains in healthy neurons co-expressing the neurofilament medium chain (NFM). a–d**: MCNs were transfected at DIV 8 with full-length NFL-FLAG (**a**), NFL(head)-FLAG (**b**), NFL(rod)-FLAG (**c**), or NFL(tail)-FLAG (**d**), together with an NFM-expressing construct. After 3 days of expression, neurons were fixed, stained with anti-FLAG and anti-NFL antibodies and the nuclear dye SYTO16, and imaged with widefield microscopy. Expanded views of regions of interest (dashed boxes) are shown inset in anti-FLAG panels. Scale bars: 20 µm (**a–d**), 10 µm (images inset in **a–d**).

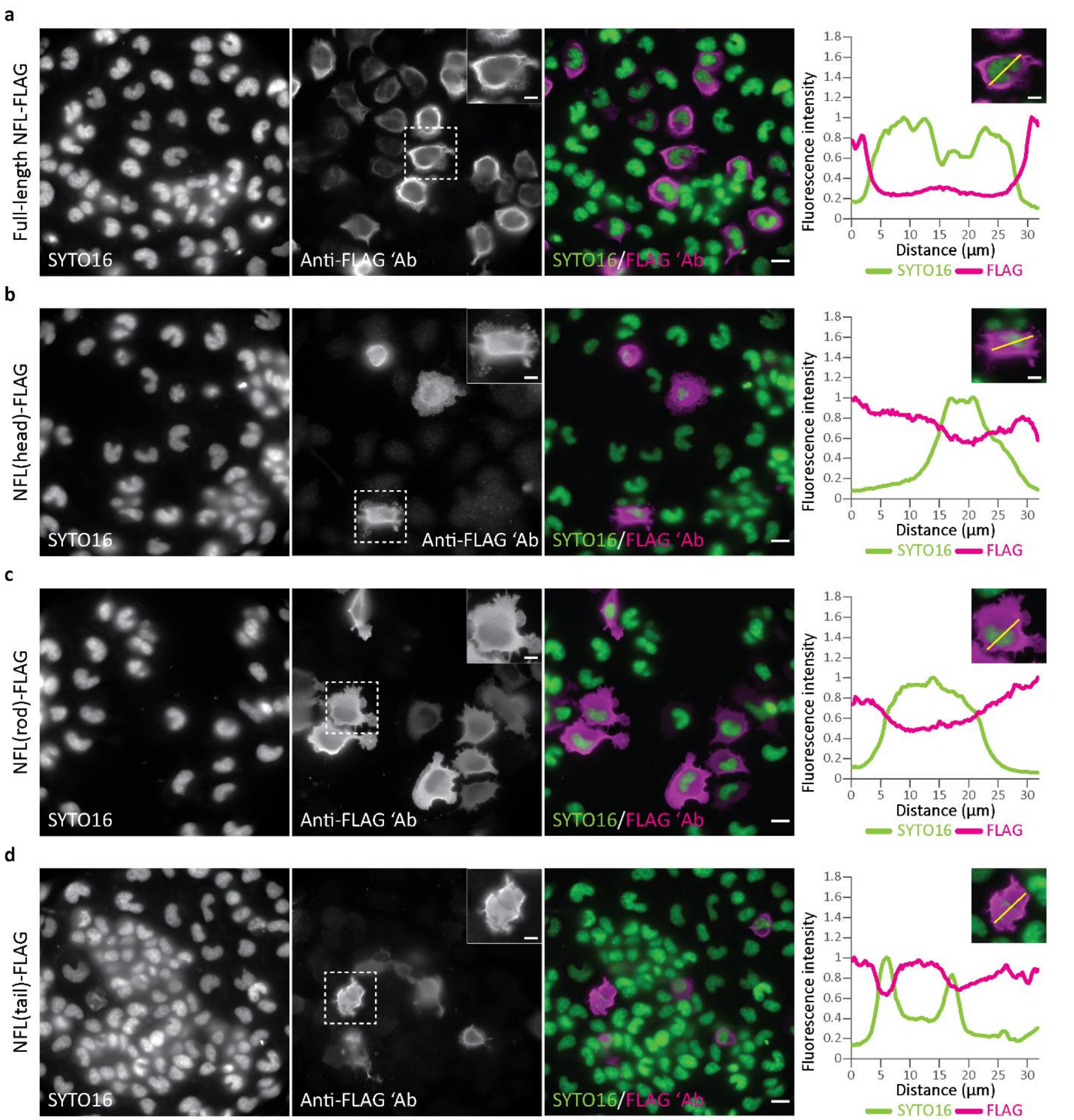


**Supplementary Fig. 8 | Localization of recombinant full-length NFL and NFL domains in healthy neuroblastoma ND7/23 cells. a–d:** ND7/23 cells were transfected with full-length NFL-FLAG (**a**), NFL(head)-FLAG (**b**), NFL(rod)-FLAG (**c**), or NFL(tail)-FLAG (**d**). On the following day, cells were fixed, stained with anti-FLAG antibody and the nuclear dye SYTO16, and imaged with widefield microscopy. Graphs on the right show line profile fluorescence intensity measurements for SYTO16 and anti-FLAG signals. Fluorescence intensities were normalized to the highest intensity value of each channel. Merged images of SYTO16 (green) and anti-FLAG (magenta) stainings (shown inset above the graphs) show the line regions of interest from which the respective fluorescence intensities were measured. Expanded views of regions of interest (dashed boxes) are shown inset in anti-FLAG panels. Scale bars: 20 µm (**a–d**), 10 µm (images inset in **a–d**).

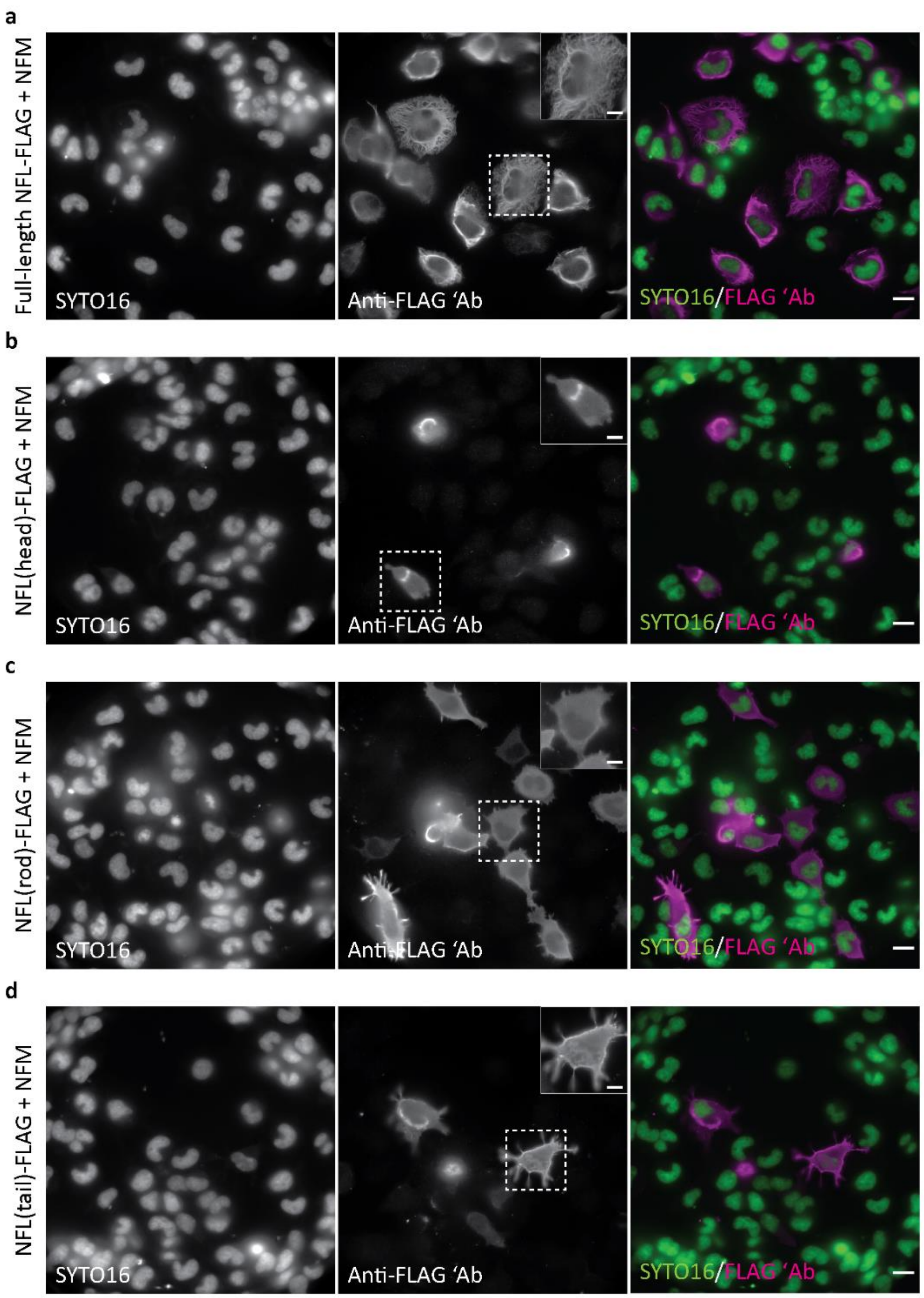


**Supplementary Fig. 9 | Localization of recombinant full-length NFL and NFL domains in healthy neuroblastoma ND7/23 cells co-expressing the neurofilament medium chain (NFM). a–d:** ND7/23 cells were transfected with full-length NFL-FLAG (**a**), NFL(head)-FLAG (**b**), NFL(rod)-FLAG (**c**), or NFL(tail)-FLAG (**d**), together with an NFM-expressing construct. On the following day, cells were fixed, stained with anti-FLAG antibody and the nuclear dye SYTO16, and imaged with widefield microscopy. Expanded views of regions of interest (dashed boxes) are shown inset in anti-FLAG panels. Scale bars: 20 µm (**a–d**), 10 µm (images inset in **a–d**).

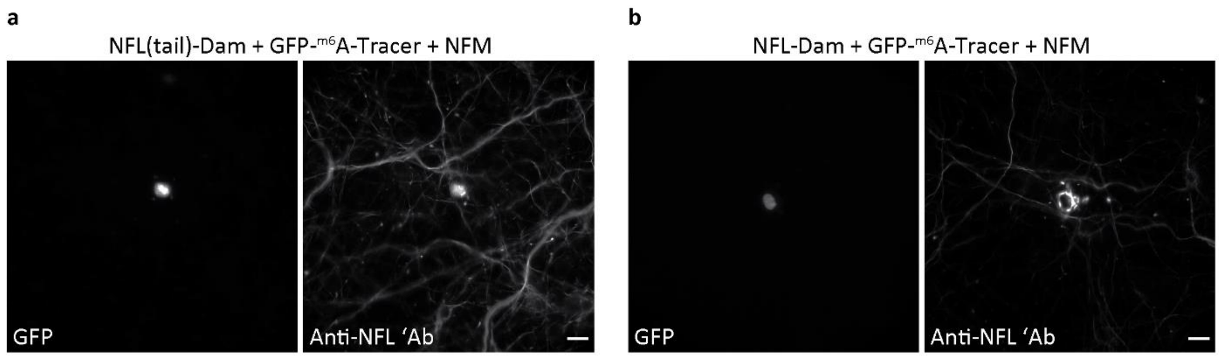


**Supplementary Fig. 10 | NFL-Dam is unable to incorporate into the neurofilament network in primary neurons.** MCNs were transfected at DIV 8 with NFL(tail)-Dam, NFM, and GFP-$^{m6}$A-Tracer (**a**) or full-length NFL-Dam, NFM, and GFP-$^{m6}$A-Tracer (**b**). On the following day, neurons were fixed, stained with anti-NFL antibody, and imaged on a widefield microscope. The brightness and contrast of GFP images were scaled linearly and in the same way in both panels so that the difference in GFP fluorescence can be readily observed. Scale bars: 20 µm.

### Supplementary Tables

**Supplementary Table 1.** Comparison of nuclear NFL fluorescence intensity by using the Kruskal–Wallis test and Dunn–Bonferroni correction for multiple comparisons

**Supplementary Table 2.** Putative calpain cleavage sites prediction in the C-terminal region of NFL and N-terminal region of GFP in the NFL-GFP construct

**Supplementary Table 3.** Primers and oligonucleotides used for cloning

**Supplementary Table 4.** Comparison of nuclear GFP fluorescence intensity by using the Kruskal–Wallis test and Dunn–Bonferroni correction for multiple comparisons

**Supplementary Table 1.** Comparison of nuclear NFL fluorescence intensity by using the Kruskal–Wallis test and Dunn–Bonferroni correction for multiple comparisons

| Groups† | Test statistic | Std. error | Std. test statistic | Significance | Adjusted significance⁺ |
|---|---|---|---|---|---|
| 1 & 2 | −773.609 | 54.294 | −14.248 | 0.000 | 0.000* |
| 1 & 3 | −536.735 | 59.498 | −9.021 | 0.000 | 0.000* |
| 1 & 4 | −501.610 | 59.186 | −8.475 | 0.000 | 0.000* |
| 1 & 5 | −759.180 | 58.093 | −13.068 | 0.000 | 0.000* |
| 1 & 6 | −520.028 | 58.885 | −8.831 | 0.000 | 0.000* |
| 1 & 7 | −388.957 | 55.650 | −6.989 | 0.000 | 0.000* |
| 1 & 8 | −337.706 | 54.230 | −6.227 | 0.000 | 0.000* |
| 2 & 3 | 236.874 | 55.906 | 4.237 | 0.000 | 0.001* |
| 2 & 4 | 271.999 | 55.574 | 4.894 | 0.000 | 0.000* |
| 2 & 5 | 14.430 | 54.409 | 0.265 | 0.791 | 1.000 |
| 2 & 6 | 253.581 | 55.253 | 4.589 | 0.000 | 0.000* |
| 2 & 7 | 384.653 | 51.792 | 7.427 | 0.000 | 0.000* |
| 2 & 8 | 435.903 | 50.263 | 8.672 | 0.000 | 0.000* |
| 3 & 4 | 35.125 | 60.668 | 0.579 | 0.563 | 1.000 |
| 3 & 5 | −222.444 | 59.602 | −3.732 | 0.000 | 0.005* |
| 3 & 6 | 16.707 | 60.374 | 0.277 | 0.782 | 1.000 |
| 3 & 7 | 147.779 | 57.224 | 2.582 | 0.010 | 0.275 |
| 3 & 8 | 199.029 | 55.844 | 3.564 | 0.000 | 0.010* |
| 4 & 5 | −257.570 | 59.291 | −4.344 | 0.000 | 0.000* |
| 4 & 6 | −18.418 | 60.067 | −0.307 | 0.759 | 1.000 |
| 4 & 7 | 112.653 | 56.899 | 1.980 | 0.048 | 1.000 |
| 4 & 8 | 163.903 | 55.512 | 2.953 | 0.003 | 0.088 |
| 5 & 6 | 239.151 | 58.990 | 4.054 | 0.000 | 0.001* |
| 5 & 7 | 370.223 | 55.761 | 6.639 | 0.000 | 0.000* |
| 5 & 8 | 421.473 | 54.345 | 7.756 | 0.000 | 0.000* |
| 6 & 7 | 131.072 | 56.586 | 2.316 | 0.021 | 0.575 |
| 6 & 8 | 182.322 | 55.190 | 3.304 | 0.001 | 0.027* |
| 7 & 8 | 51.250 | 51.725 | 0.991 | 0.322 | 1.000 |

†Groups: 1. Control; 2. nitric oxide; 3. nitric oxide + calpain inhibitor III; 4. nitric oxide + EST; 5. nitric oxide + emricasan; 6. nitric oxide + calpain inhibitor III + EST; 7. nitric oxide + calpain inhibitor III + emricasan; 8. nitric oxide + EST + emricasan.

⁺Significance values have been adjusted with the Dunn–Bonferroni correction for multiple tests.

*Indicates significance.

**Supplementary Table 2.** Putative calpain cleavage sites prediction in the C-terminal region of NFL and N-terminal region of GFP in the NFL-GFP construct

| | NFL-GFP position | Deep calpain (low threshold) | GPS-CCD (high threshold) |
|---|---|---|---|
| C-terminal region of NFL (between the antibody-recognized epitope and the C terminus) | 460 | − | + |
| | 463 | + | + |
| | 468 | + | + |
| | 469 | − | + |
| | 497 | − | + |
| | 503 | − | + |
| | 525 | − | + |
| | 531 | − | + |
| | 532 | − | + |
| | 533 | − | + |
| | 537 | + | − |
| | 538 | + | − |
| | 539 | − | + |
| N-terminal region of GFP | 545 | − | + |
| | 546 | + | − |
| | 625 | + | − |
| | 631 | + | − |
| | 633 | + | − |
| | 634 | + | − |
| | 638 | + | − |
| | 641 | + | − |

**Supplementary Table 3.** Primers and oligonucleotides used for cloning

| Purpose | Primer name | Primer sequence, 5'–3' |
|---|---|---|
| Cloning of NFL(head)-FLAG | NfL head_HindIII_fw | GGA GGA AAG CTT CACC ATG AGT TCG TTC GGC |
| | NfL head_BamHI_rv | TCC TCC GGA TCC AGC TGT GCC TTC TCT TGT GTG CG |
| Cloning of NFL(rod)-FLAG | NfL rod_HindIII_fw | GGA GGA AAG CTT CACC ATG CTG CAG GAC CTC AAC GAT CGC TTC |
| | NfL rod_BamHI_rv | TCC TCC GGA TCC AGT TCG CCT TCC AAG AGT TTT CTG TAA GCT G |
| Cloning of NFL(tail)-FLAG | NfL tail_HindIII_fw | GGA GGA AAG CTT CACC ATG GAG ACC AGG CTC AGT TTC ACC AGC |
| | NfL tail_BamHI_rv | TCC TCC GGA TCC AGA TCT TTC TTC TTA GCC ACC TGC TCC TCT C |
| Cloning of NFL(ΔA461-D543)-GFP | NFL_HindIII_fw | GCA TGC AAG CTT CAC CAT GAG TTC GTT CGG CTA CGA T |
| | NFL(A461)_BamHI_rv | TCC TCC GGA TCC GGA GCC TCA ATG GTC TCC TCG ACC TCT GTC |
| Cloning of NFL(ΔA461–D543)-FLAG | NFL_HindIII_fw | GCA TGC AAG CTT CAC CAT GAG TTC GTT CGG CTA CGA T |
| | NFL(A461)-FLAG_NotI_rv | TCC TCC GCG GCC GCT CAC TTG TCG TCA TCG TCT TTG TAG TCA GCC TCA ATG GTC TCC TCG ACC TCT GTC |
| Cloning of pcDNA3.1/Zeo(+)-Dam | HindIII_Dam_fw | GGA GGA AAG CTT ATG AAG AAA AAT CGC GCT TTT TTG AAG TGG G |
| | EcoRI_Dam_rv | TCC TCC GAA TTC TTA TTT TTT CGC GGG TGA AAC GAC TCC TG |
| Cloning of pcDNA3.1/Zeo(+)-NFL-Dam | NheI_NFL_fw | GAGGA GCT AGC CAC CAT GAG TTC GTT CGG CTA CGA TC |
| | HindIII_NFL_rv | TCC TCC AAG CTT CTT GTC GTC ATC GTC TTT GTA GTC CGG |
| Cloning of pcDNA3.1/Zeo(+)-NFL(tail)-Dam | NheI_NFL(tail)_fw | GGA GGA GCT AGC CAC CGA GAC CAG GCT CAG TTT C |
| | HindIII_NFL_rv | TCC TCC AAG CTT CTT GTC GTC ATC GTC TTT GTA GTC CGG |

**Supplementary Table 4.** Comparison of nuclear GFP fluorescence intensity by using the Kruskal–Wallis test and Dunn–Bonferroni correction for multiple comparisons

| Groups† | Test statistic | Std. error | Std. test statistic | Significance | Adjusted significance⁺ |
|---|---|---|---|---|---|
| 1 & 2 | −393.640 | 27.894 | −14.112 | 0.000 | 0.000* |
| 1 & 3 | −384.375 | 27.940 | −13.757 | 0.000 | 0.000* |
| 1 & 4 | −199.531 | 26.035 | −7.664 | <0.001 | 0.000* |
| 1 & 5 | 162.144 | 28.081 | 5.774 | <0.001 | 0.000* |
| 2 & 3 | 9.265 | 29.523 | 0.314 | 0.754 | 1.000 |
| 2 & 4 | 194.109 | 27.727 | 7.001 | <0.001 | 0.000* |
| 2 & 5 | 555.783 | 29.657 | 18.741 | 0.000 | 0.000* |
| 3 & 4 | 184.844 | 27.773 | 6.656 | <0.001 | 0.000* |
| 3 & 5 | 546.519 | 29.700 | 18.401 | 0.000 | 0.000* |
| 4 & 5 | 361.675 | 27.915 | 12.956 | 0.000 | 0.000* |

†Groups: 1. NFL-Dam; 2. NFL(tail)-Dam; 3. NFL(tail)-Dam + NFM; 4. Dam; 5. GFP-$^{m6}$A-Tracer.

⁺Significance values have been adjusted with the Dunn–Bonferroni correction for multiple tests.

*Indicates significance.
